# Covert muscle activity reveals dynamic freezing states and prepares the animal for action

**DOI:** 10.1101/2024.11.26.622641

**Authors:** Anna F. Hobbiss, Mariana C. Franco, Alexandra M. Medeiros, Charlie Rosher, César S. Mendes, Marta A. Moita

**Affiliations:** Champalimaud Research, Champalimaud Centre for the Unknown, 1400-038 Lisbon, Portugal; iNOVA4Health, NOVA Medical School|Faculdade de Ciências Médicas, NMSFCM, Universidade Nova de Lisboa, 1169-056 Lisbon, Portugal

## Abstract

When an animal detects a threat it must make a split-second choice between fight, flight or freezing (1–3). During freezing, skeletal muscles sustain tension to maintain rigid, sometimes atypical postures, for many minutes at a time (4). Meanwhile the animal must dynamically assess its surroundings to plan future actions and ready its body for movement. The interplay between the neural and somatic systems during freezing remains poorly understood. Here we show that freezing *Drosophila melanogaster* display a striking novel pattern of leg muscle activation unique to immobility, a rhythmic pulsing in the distal tibia. The muscle, which we show to be a previously undescribed leg accessory heart, displays multiple activity modes and ramps up to movement onset, implying a preparation for movement. The frequency of pulsing is dynamically modulated as the fly integrates external threat or safety cues, and artificially increasing pulse frequency leads to freezing breaks, indicating a causal role in the decision to move. Through the identification of a new *Drosophila* cardiac organ, this study provides a window into the multiple states which can underlie freezing behaviour, and the physiological changes which the body undergoes to ready the animal to move.

## Introduction

When animals are faced with a threat, a suite of defensive behaviours – fight, flight, or freezing – can be observed across the animal kingdom. The selection between these behaviours happens in a split second, and choosing the wrong one could prove fatal. Freezing, the cessation of all movement for up to minutes at a time to avoid predator detection, is common to a wide variety of phylogenetically diverse animals, from humans to invertebrates such as crayfish and *Drosophila* (5– 9). It is generally understood as a state of high alertness seen at intermediate threat levels to evade a predator’s notice and assess external surroundings, with the animal poised to perform a subsequent movement (2, 10). This idea necessarily entails a dynamic progression through the freezing state, with the animal continually integrating information from the surroundings and its own internal condition. Freezing would be maintained until the threat has increased enough to warrant a different defensive response such as fleeing, fighting, or playing dead, or decreased sufficiently to allow the resumption of other behaviours (11). The exact boundaries for the switch out of freezing should depend on various factors including internal state (such as hunger or mated state), resource availability in the environment, previous history with threats, and many more (3, 12).

A large literature on the neuronal mechanisms of defensive behaviour in vertebrates has revealed a contribution of multiple neuronal populations which show sustained activity during freezing, suggesting an active process required for freezing maintenance and other attentional aspects of the freezing state (13–17). However, strong evidence for a dynamic model of freezing is lacking, with current literature in vertebrates mainly focused on the switch between freezing and a subsequent behaviour than the progression through the freezing state itself (2, 14, 16, 18–20). Furthermore, despite being debated for more than a century, the number of different internal states which present as defensive freezing is still poorly understood (21); for instance, multiple categories have been proposed in rodents, including conditioned freezing and tonic immobility, which have varied neural implementations and may serve different purposes (22).

Until recently, the stillness of freezing led the scientific community to construe this defensive state as passive (10, 14, 16), supported by measures of cardiac activity that show a slowing down of the heart (bradycardia) during freezing in a wide range of species (23–25). However, freezing is a rapid response where atypical and non-supported poses are often adopted, meaning skeletal muscles may require a continued effort to hold a limb fixed in place against gravity (26). In rats, freezing is indeed accompanied by increased alpha motor-neuron excitability and muscle tone in the legs (4, 27) and freezing in *Drosophila* causes energy mobilization and decreased resistance to starvation, even in the presence of bradycardia (25). Additionally, lesions of the cerebellum, a brain structure responsible for fine motor control in vertebrates, cause reductions in freezing without reducing other stress responses (27), indicating that freezing behaviour requires high levels of coordination of muscles across the body. The extent to which freezing should be considered ‘passive’ or ‘active’ is therefore still strongly under debate, which is further confounded by the multiple freezing states as outlined above. Body stillness during freezing precludes any useful information being gleaned from external observation of the animal, so to further investigate if and how freezing states differ and change over time, internal state measures or proxies are necessary.

To gain insights into the neural and physiological progression of freezing, we investigated the role of a crucial somatic system, the muscles, as being the ultimate effectors of locomotor and postural control across the animal kingdom. We used *Drosophila melanogaster* as its small size and genetic manipulability make it possible to measure whole-body muscle activity in awake behaving animals to a degree that would be highly challenging in another adult animal. Crucially insect muscular systems show a high degree of conservation with vertebrate systems in terms of limb muscle organisation into antagonistic pairs contracting out of phase for limb extension or flexion, as well as the striated nature of the muscle fibres (28). As in vertebrates, muscles in insects perform not just limb movements but also visceral and circulatory functions. Pressure changes to force the flow of haemolymph (the blood equivalent) through the body are enabled by a series of muscular hearts including the main dorsal heart and various accessory hearts – independent pulsatile organs to aid haemolymph flow in structures with high energy needs, including the wings and antennae (29).

We imaged muscle activity using GCaMP in awake behaving flies, hypothesising that it would reveal a stable pattern of limb muscle activity related to the maintenance of a static posture. To our surprise, we observed instead a rhythmic pulsing of a previously unidentified muscle unit at the distal end of the fly tibia, perfectly correlated between all the 6 legs. This pulsing was a signature of immobility, which ramped up to and invariably preceded the resumption of full body movement. Different patterns of pulsing, which were responsive to threat level and external safety cues, imply that different pulsing states constitute different states of freezing. We identified the muscle as a previously unidentified accessory heart, with a crucial role in preparing the animal for subsequent movement. This unexpected finding provides a remarkably unambiguous empirical demonstration that freezing is composed of multiple internal states, and to our knowledge is the first time the activity of an arthropod heart structure has been so closely and causally linked with the onset of movement.

## Results

We imaged the muscle activity of behaving *Drosophila melanogaster* using the calcium sensor GCaMP6f under the control of myosin heavy chain muscle driver (mhc-LexA) as proxy for muscle contraction (30). Flies were tethered by the thorax and given a small polystyrene ball, upon which they could perform various behaviours including walking, grooming and egg laying (25). To promote robust walking, a dark vertical stripe (buridan stripe (31)) was presented throughout the experiment on a frontal monitor (Fig. 1A). To induce freezing behaviour, a threat in the form of a looming stimulus – an expanding black disc on a lighter background, which mimics an object on collision course (32) – was presented on the monitor. After a 5-minute baseline period 96% of flies froze to the onset of the first loom (Fig. 1B, Fig. S1A). Subsequent looms were presented once the fly had broken from freezing to resume walking or grooming (see Methods for details).

**Fig. 1.**
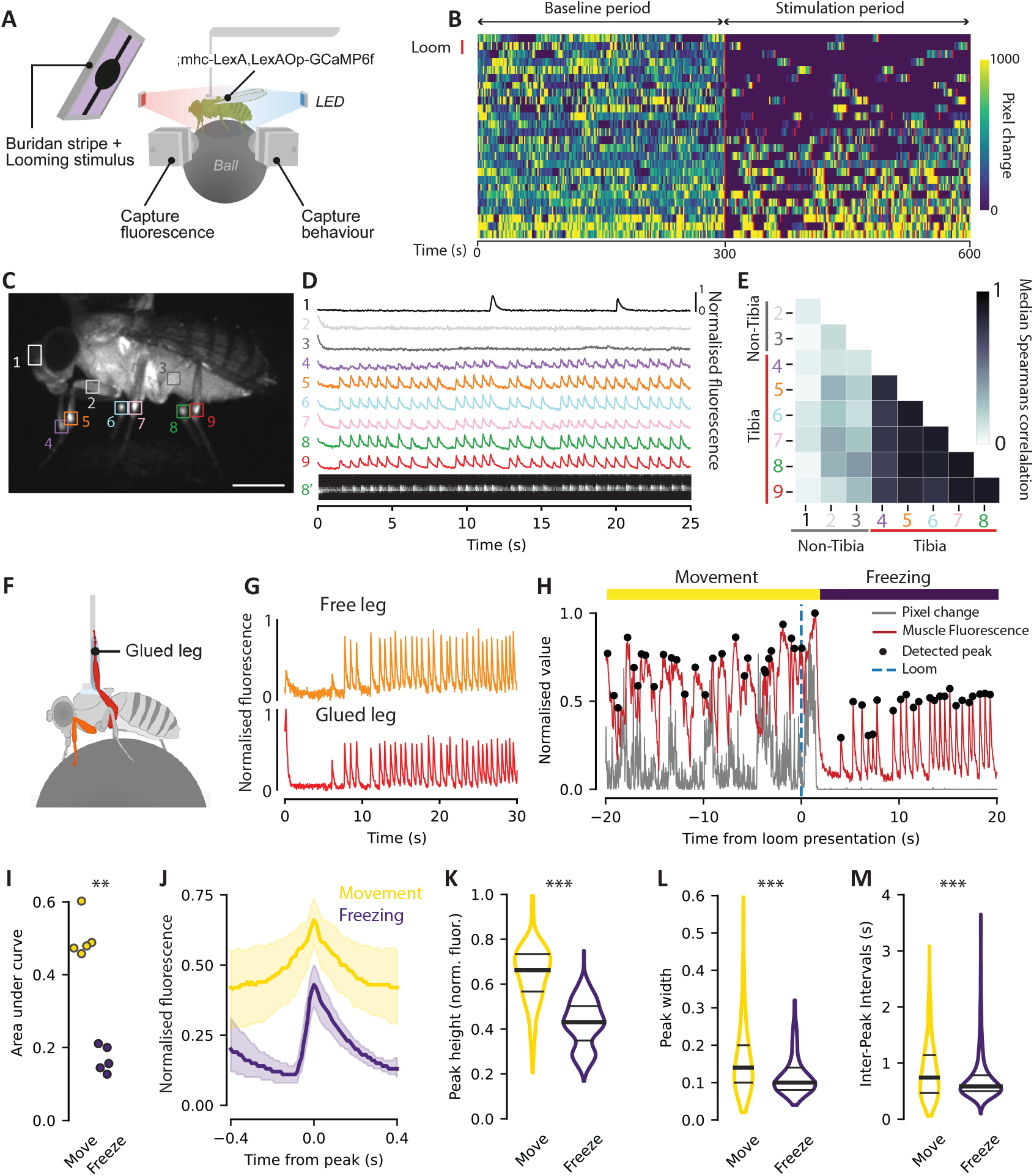
*Drosophila* display pulsing in a distal tibia muscle during immobility. A) Schematic of tethered fly behavioural set-up. B) Pixel change over the experiment (cropped at 600 s). Looming stimuli are displayed as red lines. Each row corresponds to a fly, rank ordered by the amount of time spent freezing during the stimulation period. *N* = 26 flies. C) Example of GCaMP6f signal in a tethered fly, showing locations of the 9 ROIs used for analysis in D and E. Scale bar = 500 µm. D) Example trace of normalised fluorescence changes in 9 ROIs during a freezing bout (1-9). The bottom trace shows a fluorescence line profile for ROI 8. E) Correlation matrix for moment-to-moment fluorescence correlations between the 9 ROIs. *N* = 15 immobility bouts (11 flies). F) Schematic of glued-leg preparation. The hind glued leg is shown in red, and a free leg used as a comparison in orange. G) Example of fluorescence changes in distal tibial muscles of a free leg (orange) and the glued leg (red) during a freezing bout. H) Example of glued leg fluorescence before and after freezing onset. Y axis spans the minimum (0) and maximum (1) values for the experiment for both pixel change (plotted in grey) and muscle fluorescence (plotted in red). I) Integration of area under fluorescence intensity curve per second, for movement vs freezing bouts during the stimulation period. *N* = 5 flies. *P* = 0.008, Mann Whitney U test. J) Average peak shape (median and interquartile range) for movement vs freezing bouts. *N* Movement = 489, *N* Freezing = 1344 (5 flies). K) Normalised height of fluorescence peaks during stimulation period. *N* Movement = 489, *N* Freezing = 1344 (5 flies). *P* < 0.001, Mann Whitney U test. L) Width of fluorescence peaks during stimulation period. *N* Movement = 494, *N* Freezing = 1344. *P* < 0.001, Mann Whitney U test. M) Inter-peak intervals during stimulation period. *N* Movement = 467, *N* Freezing = 1316, *P* < 0.001, Mann Whitney U test.

### Distinct leg muscle pulsing is observed during immobility

We predicted that imaging muscle activation using GCaMP would reveal a stable pattern of limb muscle activity related to the maintenance of a static posture – either quiescence (33), or a prolonged activation of the muscles (34, 35). Although posture is held rigidly during freezing behaviour, as evidenced by the lack of pixel change throughout freezing bouts (see Methods), muscle activity was surprisingly not constant. Instead, a salient, rhythmic muscle signal was observed in the distal end of the tibia in all flies (Regions Of Interest (ROIs) 4-9 in Fig. 1C and D). This signal was strikingly correlated between all the legs of the fly (Fig. 1D and E, Fig. S1B, Supp Video 1). Calcium signals could be seen in other ROIs, such as the head, but these were not pulsatile or correlated with those observed in the distal tibia (Fig. 1D and E, Fig. S1B). Although this signal was most salient during the long, loom-induced freezing bouts, it could also be observed in moments of short spontaneous immobility in the baseline (Fig. S1C), and in immobile legs during grooming (Fig. S1D). To ensure that the pulsing activity was not an artefact of the tethering procedure or gripping the ball, we imaged freely moving flies and saw moments of correlated pulsing activity during grooming and freezing (Fig. S1E, Supp Video 2).

A correlated muscle activity between all legs was surprising since during normal movement the muscles of different legs work in an out-of-phase fashion. We asked if the pulsing signal was specific to moments of immobility, or if it continued throughout movement. It is technically challenging to image muscle activity of moving legs. If correlated pulsing activity was maintained throughout movement however, imaging from a single immobilised leg, whose activity would faithfully follow that of the other five free legs, could be used as a proxy for muscle activity during movement. We immobilised a single leg by gluing it vertically to the tethering wire and performed our looming protocol while the fly walked on a ball, allowing us to assess distal tibia muscle activity throughout the whole experiment in the immobilised leg (Fig. 1F). During moments of immobility pulsing was correlated between the glued leg and free legs (Fig. 1G), validating our experimental approach. We then compared muscle activity in the glued leg during different behaviours. During movement, fluorescence was elevated and showed a more variable profile compared to when the fly was immobile, when stereotyped pulses were visible (Fig. 1H and I, *P* = 0.008). To better describe the fluorescence dynamics, we detected peaks throughout the experiment and calculated a peak-triggered-average of fluorescence for the two behavioural states (Fig. 1J). During freezing, the peak shape was sharp and largely invariant reaching a mid-level fluorescence, while peaks during movement were wider and more variable (Fig. 1J-L, *P* < 0.001). Inter-peak intervals were both shorter and less variable during freezing (Fig. 1M, *P* < 0.001), implying a more stereotyped activity. The same results were also found for non-loom-triggered (spontaneous) immobility vs movement during the baseline, albeit with a higher variance since extended periods of spontaneous immobility were rare (Fig. S1F-J). These data indicate that distal tibial muscles in flies show a signature pulsing signal during moments of immobility, distinct from their activity during movement.

### The pulsing muscle is an accessory heart

We next sought to identify the pulsing muscle to understand its biomechanical features and infer its functional role. The *Drosophila* leg consists of 9 segments: the coxa, the trochanter, the femur, the tibia, and 5 tarsal segments (Fig. 2A). Previous descriptions of muscles in the legs defined 14 muscles per fly leg, consisting of one or more flexor, extensor, and reductor muscles per segment which control the position of the subsequent limb segment, as well as a long tendon muscle in the femur and the tibia to control the distal tip of the tarsus (36, 37) (see Methods for details of our usage of muscle nomenclature). Since none of these descriptions captured the apparent location, small size, and independent muscle activity we observed, we used spinning disc confocal microscopy to image an immobilised leg of a live fly expressing muscular GCaMP, glued on its side to a cover slip (Fig. S2A). Muscle activity of coactive muscle groups could be clearly distinguished through the cuticle (Fig. 2B, Supp Video 3). We observed a subset of muscle fibres at the distal end of the tarsus flexor muscle with a characteristic rhythmic pulsing activity and extracted the mean fluorescence levels from this ROI (termed ‘1a’), from an ROI in a more proximal section of the tarsus flexor muscle (‘1b’) and another 3 ROIs drawn inside leg muscles (Fig. 2B and C) (see Methods for muscle nomenclature). We classified muscle activity into 4 states: sustained plateaus of high fluorescence (‘High’); quiescent moments with no fluorescence (‘Low’); rhythmic pulsing reaching intermediate fluorescence levels (‘Pulsing’); and transitional states which did not fit any of these criteria (‘Transition’) (Fig. S2B). Whilst high and low states were observed in all muscle groups, pulsing was highly enriched in ROI 1a (with a small increase also seen in 1b) but was almost absent in other muscle groups (Fig. 2D).

**Fig. 2.**
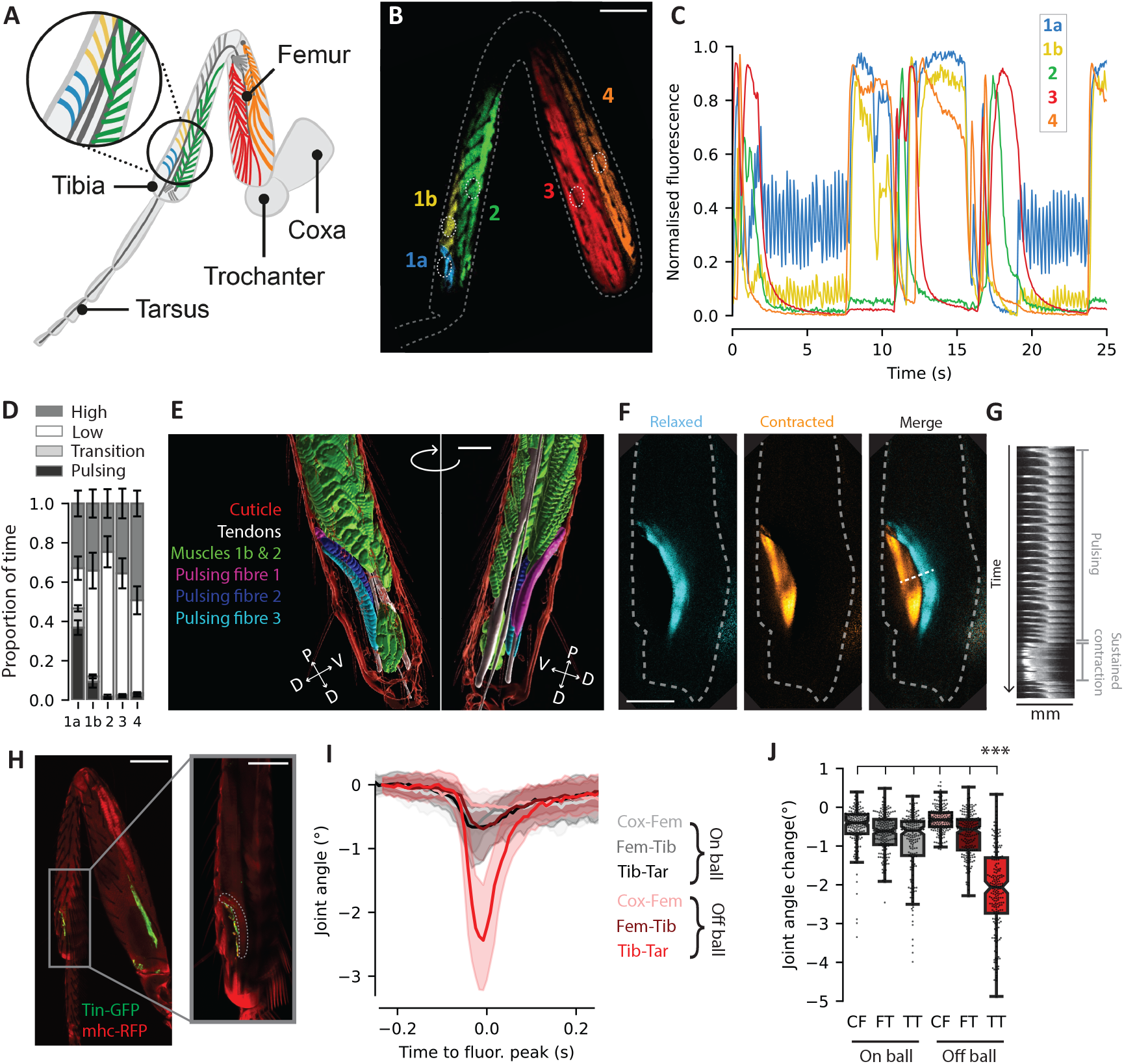
The distal fibres of the tarsus flexor muscle form an accessory heart. A) Schematic of a *Drosophila* leg. Muscle colours represent regions which are measured in parts B-D. Blue: Distal fibres of tarsus flexor muscle; Yellow: proximal fibres of tarsus flexor muscle; Green: tarsus extensor muscle; Red: tibia flexor muscle; Orange: Tibia extensor muscle. Scale bar = 100 µm B) False colour confocal image showing co-active regions which were measured using the labelled ROIs (white dotted lines). Colours correspond to muscles delineated in (A). C) Individual example of normalised fluorescence values of muscles in an immobilised leg. *N* = 9 flies. D) Stacked bar charts of overall proportion of time spent in different activity states for different muscles. Error bars = SEM. E) 3D reconstruction of the distal tibia, with the 3 fibres of the pulsing muscle (1a) separated by colour. Scale bar = 40 µm. F) Spinning disc images of pulsing fibres (1a) labelled with a membrane-bound GFP in a relaxed (cyan) and contracted (orange) state. Scale bar = 30 µm. White dashed line was used to create the fluorescence profile in (G). G) Fluorescence line profile drawn perpendicular to the pulsing fibres. Moments of pulsing and sustained contraction are indicated. H) Confocal image of Tin-Gal4-GFP (green) and mhc-RFP (red). Note there is strong autofluorescence of the cuticle in the red channel. Scale bar = 100 µm. Inset zoom in on the indicated region of the distal tibia with pulsing fibres outlined with dotted white line. Scale bar = 50 µm. I) Leg angle displacement (median and interquartile range) for legs which are attached to the substrate (On ball) and not attached (Off ball), centred around the time of a fluorescence pulse. *N* On ball = 177 pulses (3 flies). *N* Off ball = 205 pulses (3 flies). J) Joint angle change for the Coxa-Femur, Femur-Tibia and Tibia-Tarsus joints around the moment of pulse occurrence. *N* On ball = 177 pulses (3 flies). *N* Off ball = 205 pulses (3 flies). *P* Off ball TT < 0.001 as compared to all other values (Dunn’s test).

Pulsing in 1a was seen at the expense of the quiescent state and was concomitant with the low state in all other ROIs (Fig. 2C and D, Fig. S2C). In addition, as expected for a canonical antagonistic coupling of muscle activation as would be observed during walking, ROI 1a showed predominantly high and low states together with ROIs 1b and 4, whose contraction would raise the leg, and in anti-correlation with ROIs 2 and 3, which would lower the leg (Fig. S2C). These results are consistent with the high/low states corresponding to attempted movement and the pulsing state as signature of immobility. Our results indicate a partial dissociation in the activity of ROIs 1a and 1b, both in the tarsus flexor muscle, with the distal-most fibres (1a) active most of the time, either in the high or pulsing state, while the more proximal fibres (1b) stay either quiescent or, when active, show a predominance of the high activity state. A momentto-moment correlation of fluorescence levels reveals that during pulsing, 1a activity dominates with little contribution of 1b, but when 1a enters a state of strong sustained contraction, 1b is also co-opted and contracts strongly (Fig. S2D, Supp Video 4).

A functional dissociation of the distal fibres of the tarsus flexor muscle is further strengthened by our discovery that these fibres share a distinct genomic signature not expressed by the other muscles in the leg, as shown by their labelling by the Gal4 line CLIP-190 (38) (Fig. S2E). Using this evidence, we used a manually segmented confocal image of leg, tendon and cuticle structures in the distal tibia to make a 3D reconstruction in Imaris (Fig. 2E, Supp Video 5). We distinguished the pulsing fibres from the other leg muscles using their attachment points which are distinct from the rest of the tarsus flexor muscle. This revealed that the pulsing muscle consists of 3 fibres which overlay the tarsus flexor tendon and form a distinctive curved bow shape around a muscle-free pocket or chamber in the dorsal side of the distal tibia. When flies expressing GFP fused to tropomyosin (an essential component of actin strands of striated muscle) were injected with the soluble blue dye erioglaucine, which diffused around the body freely, the dye accumulated inside this chamber indicating that it is filled with haemolymph (Fig S2F). This structure was reminiscent of the enlarged haemolymph-filled sinuses described in other insect species associated with accessory hearts – pulsatile structures which aid in haemolymph flow through body appendages by creating oscillating pressure changes (39, 40). Indeed, live imaging of leg muscle fibres using spinning disc microscopy revealed that, unlike all other muscles, 1a muscle contraction caused a significant shape change. In its relaxed state, the fibres of muscle 1a formed a curved bow; however, as the muscle contracted, it moved towards the dorsal cuticle into the adjacent pocket. At its maximum contraction, it became completely straight with no curve, reducing the volume of the pocket (Fig. 2F and G, Supp Video 6). Reduction of the space inside the pocket would cause pressure changes to the haemolymph inside it. These data therefore suggest that muscle pulsing can act as an accessory heart, causing changes to pressure inside the leg to propel haemolymph along it. Haemolymph flow is important for nutrition and turgidity, which could be crucial for posture and movement onset. This would also explain why the pulsing is exclusively a phenomenon observed during immobility, as during normal movement contractions of the complete tarsus flexor muscle set (1a and 1b) would cause pressure changes as a by-product of movement, rendering additional pulsing activity unnecessary. To confirm this intuition, we imaged cells expressing tinman (41), a genetic marker of heart cells, using a tin-Gal4 driver (42) and found it to be expressed on the distal surface of the 1a fibres, reinforcing their role as an accessory heart (Fig. 2H). This is to our knowledge the first description of a leg accessory heart in *Drosophila* or indeed in Diptera (thus far leg hearts have only been reported in locusts and Hemiptera) (40).

Given the attachment of the muscle 1a fibres to the tarsus flexor tendon (Fig. 2E), they could still influence leg movement in addition to their role as an accessory heart. We tracked fly leg segments using DeepLabCut (43) to quantify joint angle changes associated with pulsing. During normal freezing, when the fly was holding the ball, there was negligible movement. However, on rare occasions when the fly dropped the ball but continued freezing, a small but obvious twitch of the tarsi, resulting from a reduction in the angle between the tibia and the tarsus (the expected movement from pressure applied to the tarsus flexor tendon) was observed concurrently with pulsing (Fig. 2I and J, Supp Video 7). Movement deflection was negligible at other joints (*P* < 0.001). The muscle thus remains capable of exerting force, as would be expected if it still participates in overt tarsal movements as we show in Fig. 1F-M. The lack of movement associated to muscle pulsing when legs are in braced on a ball lends weight to the hypothesis that its main function is that of an accessory heart, whose pulsing activity during immobility aids haemolymph flow and thus the turgidity and nutrition of leg structures. In published and anecdotal evidence, similar correlated leg twitches have been observed in various different preparations when the fly is immobile, including anaesthesia recovery and sleep (44), which we can now confidently assign to the activity of the tarsal leg heart.

### Distinct pulsing patterns during freezing show ramping towards movement resumption

We next sought to understand the patterns of muscle activation and their relation to freezing dynamics. For our original experiments in which tethered flies froze to a loom (as shown in Fig. 1B) we examined the patterns of pulsing within loom induced freeze bouts, as well as for the rare self-initiated pauses during the baseline (Fig. 3A). The extent to which flies pulsed was very variable, with frequencies changing over time. In both baseline and stimulation conditions, flies progressed from a non-pulsing to a pulsing state before breaking from freezing, with the frequency of pulsing ramping up to the moment of movement onset (Fig. 3B and C, *P* = 0.001). Notably, a fly breaking from freezing and commencing a full-body movement was never seen without at least one preceding pulse (see Methods for details). Although pulsing seemed to inevitably precede the start of a movement bout, the patterns of pulsing within freezing bouts were diverse. Notably, some freezing bout contained very long periods of time with no pulses (up to 80s), which were not observed in spontaneous baseline pauses which tended to be much shorter (Fig. 3A). To examine different patterns of pulsing during freezing we designed a classifier to separate the distribution into two groups (Fig. S3A, see also Methods for criteria and quantification). We broadly characterised the groups as *Continuous* (Fig. 3D and Fig. S3B), during which the pulsing occurs throughout the majority of the freezing bout, and *Delayed* (Fig. 3E and Fig. S3C), whereby there is a prolonged length of time without pulsing at the beginning of the bout. We then asked whether there was any behavioural relevance to the observed differences in the pulsing of the leg heart. Indeed, pulsing dynamics (even after excluding the initial long pause in the Delayed bouts) were different for the two conditions, being both faster and more regular during Continuous bouts than for Delayed (Fig. 3F-H, *P* < 0.001). In both cases, as we had seen for the pooled data, pulse frequency ramped up to movement onset (Fig. 3I, Fig. S3D and E, *P* = 0.001). We then looked at the length of the immobility bouts and saw that the Delayed bouts were on average longer than Continuous bouts (Fig. 3J, *P* = 0.015). The necessity for pulsing before movement onset, the ramping up of the frequency before movement commenced, and the increased length of time of Delayed bouts, strongly suggest that pulsing represents a state of preparedness to move.

**Fig. 3.**
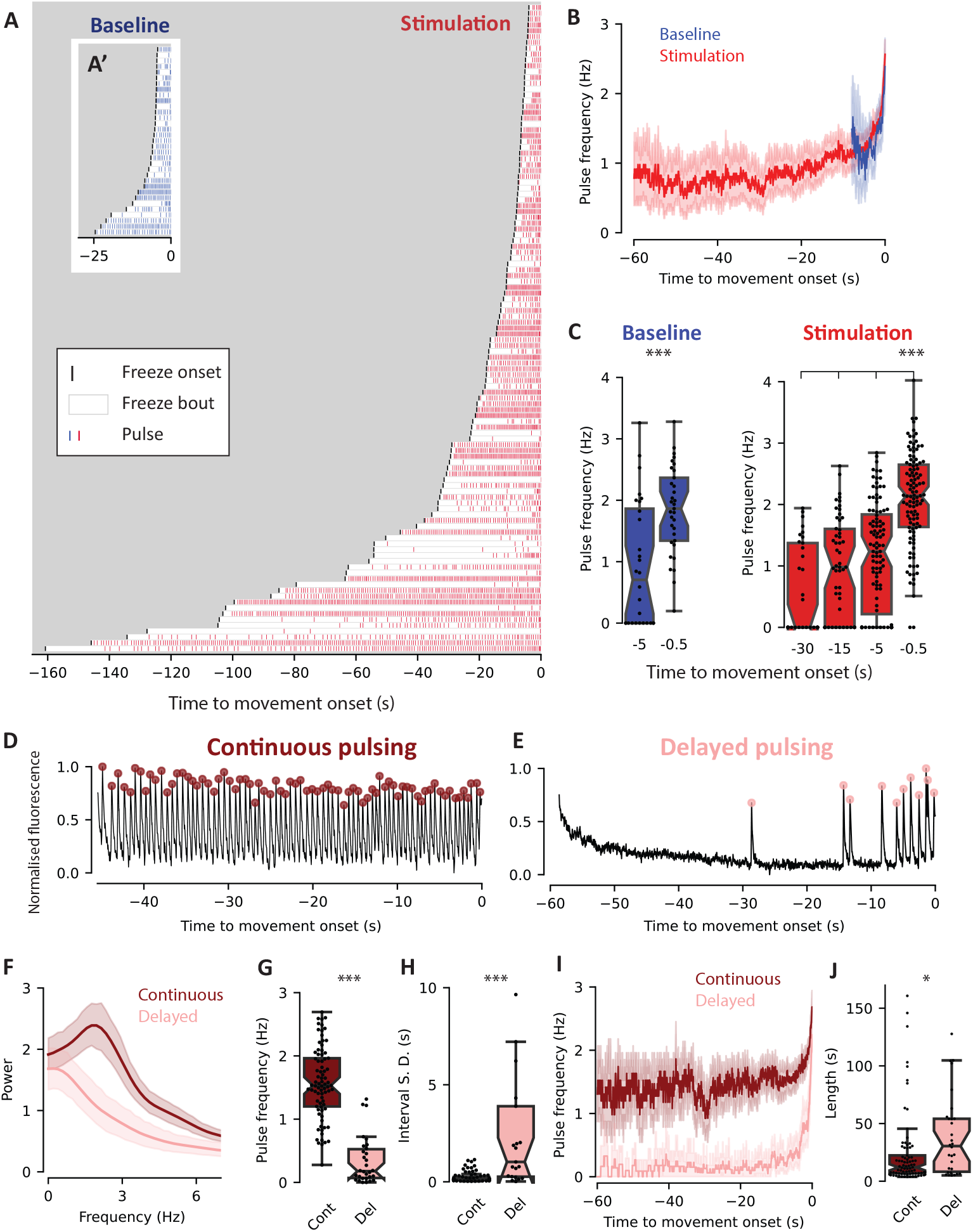
Pulsing shows heterogenous states and precedes movement onset. A) Freezing bouts longer than 4 seconds, showing moments of muscle pulsing. Bouts are aligned to movement onset and ordered by length of freezing bout. *N* = 111 bouts (26 flies). A’) As in A but showing spontaneous immobility bouts from the baseline period. *N* = 33 bouts (26 flies). B) Pulse frequency of immobility/freezing bouts for baseline/stimulation periods respectively, aligned to the moment of movement onset. *N* Baseline = 33 bouts, *N* Stimulation = 111 bouts (26 flies). C) Indicated timepoints of pulse frequencies for baseline and stimulation periods. *N* Baseline = 26 (−5 s), 33 (−0.5 s), *P* = 0.001. *N* Stimulaton = 33 (−30 s), 54 (−15 s), 104 (−5 s), 111 (−0.5 s), (26 flies), *P* = 0.001 (0.5 s as compared to all other timepoints) (Nemenyi-Friedman test). D) Normalised fluorescence in pulsing muscle in an example of Continuous pulsing. Dark red: detected peaks. E) As in (D) for Delayed pulsing. Light red: detected peaks. F) Power Spectral Density of pulsing extracted from fluorescence signals for Delayed vs Continuous bouts. *N* Continuous = 81, *N* Delayed = 28 (26 flies). G) Frequency of pulsing bouts for Continuous and Delayed. *N* Continuous = 81, *N* Delayed = 30, (26 flies), *P* < 0.001 (Mann Whitney U test). H) Standard deviation of inter-pulse interval for Continuous vs Delayed. *N* Continuous = 80, *N* Delayed = 21 (26 flies), *P* < 0.001 (Mann Whitney U test). I) Pulse frequency of freezing bouts for Continuous or Delayed bouts, aligned to the moment of movement onset. *N* Continuous = 81, *N* Delayed = 30 (26 flies). J) Length of freezing bouts for Continuous or Delayed. *N* Continuous = 81, *N* Delayed = 30 (26 flies). *P* = 0.015.

### Pulsing state is dynamically modulated by external threat intensity and safety cues

An implication of our model is that different pulse dynamics represent different states of freezing, with Pulsing being a state of preparedness to move, and Delayed bouts representing a deeper state of immobility. These differences could be simply due to interanimal variability (a pulsing ‘personality’) or could represent flexible choices between different freezing states. For canonical defensive behaviours such as fleeing and freezing, the strength of the threat impacts the decision of the animal, biasing the choice to one behaviour or another (45, 46). We sought to test whether changing threat strength affected the pulse type displayed. We adapted our visual stimulus to increase the saliency of the threat by decreasing the contrast of the occluding buridan stripe (Fig. 4A), leading to a 20% contrast difference between the stripe and the loom, which we called High Saliency. We then compared the behaviour and pulsing of flies during freezing induced by the High Saliency and the previous Low Saliency (0% contrast difference) stimuli, as used in Fig. 1 and 3. Changing the buridan stripe contrast did not change baseline behaviour (Fig. S4A); however, increasing the threat saliency strongly biased the freezing type towards Delayed pulsing (Fig. 4B, Fig. S4B, *P* < 0.001). When we compared the profiles of individual flies, we saw that in both cases most flies showed a strong preponderance to a single response type (either Continuous or Delayed), while a smaller subset of flies showed a mixture of Continuous and Delayed pulsing (Fig. S4C and D). The modulation of freezing state by external clues shows that the state the animal adopts is flexible and not simply an intrinsic, stereotyped response of individual flies, and the shift towards Delayed pulsing at higher threat levels implies that Delayed pulsing represents a deeper freezing state than Continuous.

**Fig. 4.**
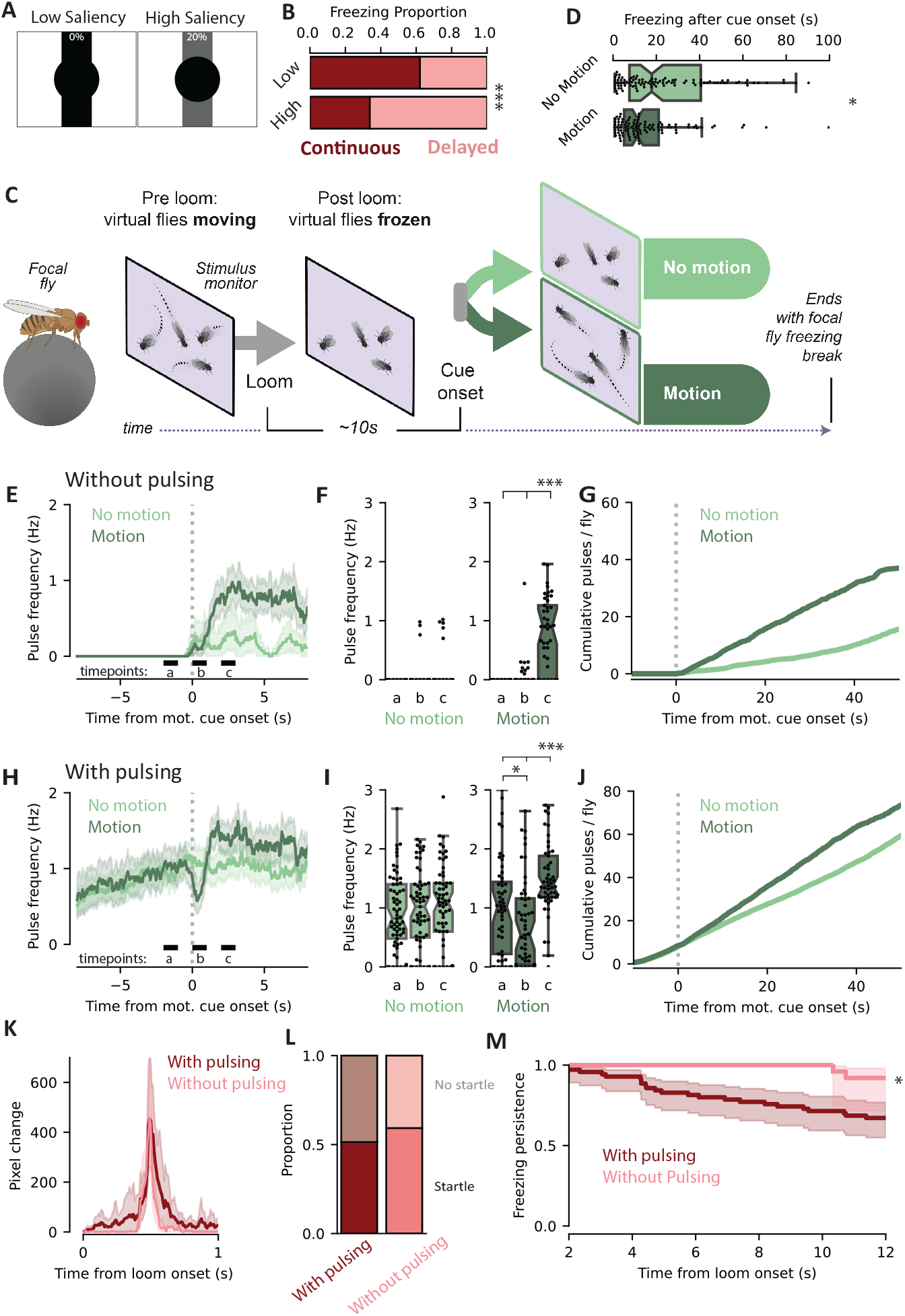
Pulsing state is modulated by threat valence and external information. A) Schematic of Low Saliency and High Saliency looming stimuli. B) Proportion of freezing time spent in Continuous or Delayed mode, for Low and High Saliency stimuli. *N* Low Contrast = 25 flies, *N* High Contrast = 20 flies, *P* < 0.001, Chi-squared test. C) Experimental protocol for virtual social motion stimulus. D) Length of time spent freezing after cue onset. *N* No Motion = 102 bouts (33 flies), *N* Motion = 103 bouts (35 flies), *P* = 0.026, Mann Whitney U test. E) Pulse frequency for the time around the onset of the motion cue, for flies that were Without Pulsing before cue onset. ‘a’, ‘b’ and ‘c’ relate to the timepoints used for quantification in (F). *N* No motion = 29 bouts, *N* Motion = 48 bouts. F) Pulse frequency for the timepoints indicated in (E), for flies that were Without Pulsing before cue onset. No Motion: *N* = 28 bouts, *P* a-b = 0.84, *P* a-c = 0.53, *P* b-c = 0.84. Motion: *N* = 45 bouts, *P* a-b = 0.54, *P* a-c = 0.001, *P* b-c = 0.001, Friedman test. G) Cumulative frequency of pulses per fly, for flies that were Without Pulsing before cue onset. *N* No Motion = 28 bouts, *N* Motion = 45 bouts. H) As in (E), for flies that were With Pulsing before cue onset. *N* No motion = 73 bouts, *N* Motion = 55 bouts. I) As in (F), for flies that were With Pulsing before cue onset. No Motion: *N* = 65 bouts, *P* a-b = 0.25, *P* a-c = 0.09, *P* b-c = 0.8. Motion: *N* = 53 bouts, *P* a-b = 0.046, *P* a-c = 0.001, *P* b-c = 0.001, Friedman test. J) As in (G), for flies that were With Pulsing before cue onset. *N* No Motion = 65 bouts, *N* Motion = 53 bouts. K) Pixel change after the loom for flies freezing at loom onset. *N* With Pulsing = 72 bouts, *N* Without Pulsing = 27 bouts. L) Proportion of freezing flies showing a startle response. *P* = 0.6, Chi-squared test. *N* With Pulsing = 28 flies, *N* Without Pulsing = 28 flies. M) Freezing bout persistence for Without Pulsing vs With Pulsing flies. *P* = 0.016, Kaplan-Meier estimator.

Even in a Delayed state, pulsing was always initiated before movement onset. We wondered what controlled the transition from quiescence to pulsing, and whether it was influenced by the external environment. It was recently shown that visual social motion from the movement from surrounding flies can act as a safety cue to a freezing fly and promote freezing breaks (12). We designed a virtual ‘social motion’ cue by displaying a video of walking flies on the stimulus monitor in front of our focal tethered fly (Fig. 4C). The virtual flies moved continuously throughout the baseline period. Upon loom presentation, the video was paused (giving the appearance of the virtual flies freezing) for a period of ∼10 seconds. Flies could then be presented with one of two cues: either ‘No Motion’, where the video stayed paused, or ‘Motion’, where the video was restarted causing the virtual flies to start moving and presenting the social safety cue to the focal fly (see Methods for full description and rationale). We observed that, in line with experiments in freely moving flies surrounded by real conspecifics, the focal fly broke from freezing more quickly when exposed to the motion cue (Fig. 4D, *P* = 0.026).

We next examined the pulsing characteristics during freezing bouts. We divided the flies into two groups based on the number of pulses in the 10 seconds before the cue onset: ‘Without pulsing’ (0 pulses) and ‘With pulsing’ (≥ 1 pulse) (Fig. S4E and F). In the ‘Without pulsing’ group, a marked upregulation of pulse rate was visible in the Motion condition compared to the No Motion, starting 1-2 seconds after the onset of social motion (Fig. 4E-G, *P* = 0.001), showing that although the overt behaviour of the flies (freezing) is not altered, their somatic state is modulated by the cue. In the ‘With pulsing’ group, we saw a sharp decrease in the pulsing rate immediately following the onset of the social motion cue, followed by an upregulation (Fig. 4H-J, *P* = 0.001). The initial drop was interestingly reminiscent of the transient cardiac arrest – the proverbial ‘heart skipping a beat’ – upon a sudden stimulus observed in many animals including *Drosophila* (25), suggesting that the immediate onset of the motion cue was unexpected; however subsequently a similar upregulation as in the ‘Without pulsing’ group was observed. These experiments suggest that a safety signal, the motion cue, leads the flies to upregulate their pulsing rate in preparation for movement onset, indicating that environmental changes do influence the flies’ somatic state.

Our previous data suggest that pulsing invariably preceded voluntary non-cued movement onset. We asked if flies in a non-pulsing state were physically unable to break from freezing even in the face of an external cue. We adapted our original stimulus protocol to deliver looms on an automatic schedule with an average interval of 15s. This would result in another threat stimulus being delivered to currently freezing flies, which can lead to freezing breaks (9) (Fig. S4G). We examined the instances when a loom was delivered whilst the fly was freezing and divided them based on their pulsing state in the 5 seconds before the loom. We chose two groups belonging to the extremes of the distribution – ‘Without pulsing’ (those which showed no pulses) and ‘With pulsing’ (those which showed at least three pulses). When we plotted the pixel change, a proxy for movement, immediately after the onset of the loom, a large proportion of the flies in both conditions showed a sudden startle response to the loom as shown by a sharp peak in pixel change (51.3% for pulsing, 59.2% for quiescent, (Fig. 4K and L). This demonstrates that flies are not paralysed in a non-pulsing state and could startle to an external cue in a very similar manner to those in a pulsing state, with no discernible differences in response onset or magnitude. A startle response is however a qualitatively different behaviour than the coordinated, voluntary movement initiation which we studied in Figs 1 and 3, and indeed almost all flies tested returned to freezing immediately after the startle response (Fig. 4M). We asked whether the pulsing state of the fly prior to threat delivery affected the subsequent likelihood of a fly to break from freezing in a sustained manner. We plotted survival curves of the freezing bout length after the loom delivery and saw that flies which had previously shown pulsing had reduced freezing durations as compared to the quiescent flies (Fig. 4M, *P* = 0.016). These findings imply that whilst a fly in a quiescent state is physically able to engage the motor system if forced to by external circumstance (e.g. to startle to another threat), it is likely to then revert to the previous extended freezing state, whereas a fly continuously pulsing is more likely to resume walking.

### Freezing neural circuitry modulates pulsing rate

We next wondered how known circuits which enact freezing states in flies modulate pulsing state. We leveraged a pair of descending neurons, DNp09, which have been shown to be vital for freezing behaviour (9). We used flies expressing GCaMP in the muscles driven by mhc-LexA, and the red-sensitive channelrhodopsin Chrimson in DNp09 neurons driven by DNp09-Gal4 to allow activation of these neurons upon red light presentation. We imaged flies in the spinning disc set-up and analysed them using the same ROIs as described in Figure 2. When DNp09 neurons were optogenetically activated with a constant 10 second red-light pulse, fluorescence in all leg muscles immediately decayed except for pulsing activity in ROI 1a (and to a lower level in ROI 1b), which continued throughout the stimulation period (Fig. 5A, Fig. S5A). This indicated that, similarly to loom-induced freezing, DNp09-induced freezing could silence muscles needed for movement whilst pulsing activity in ROI 1a was left intact.

**Fig. 5.**
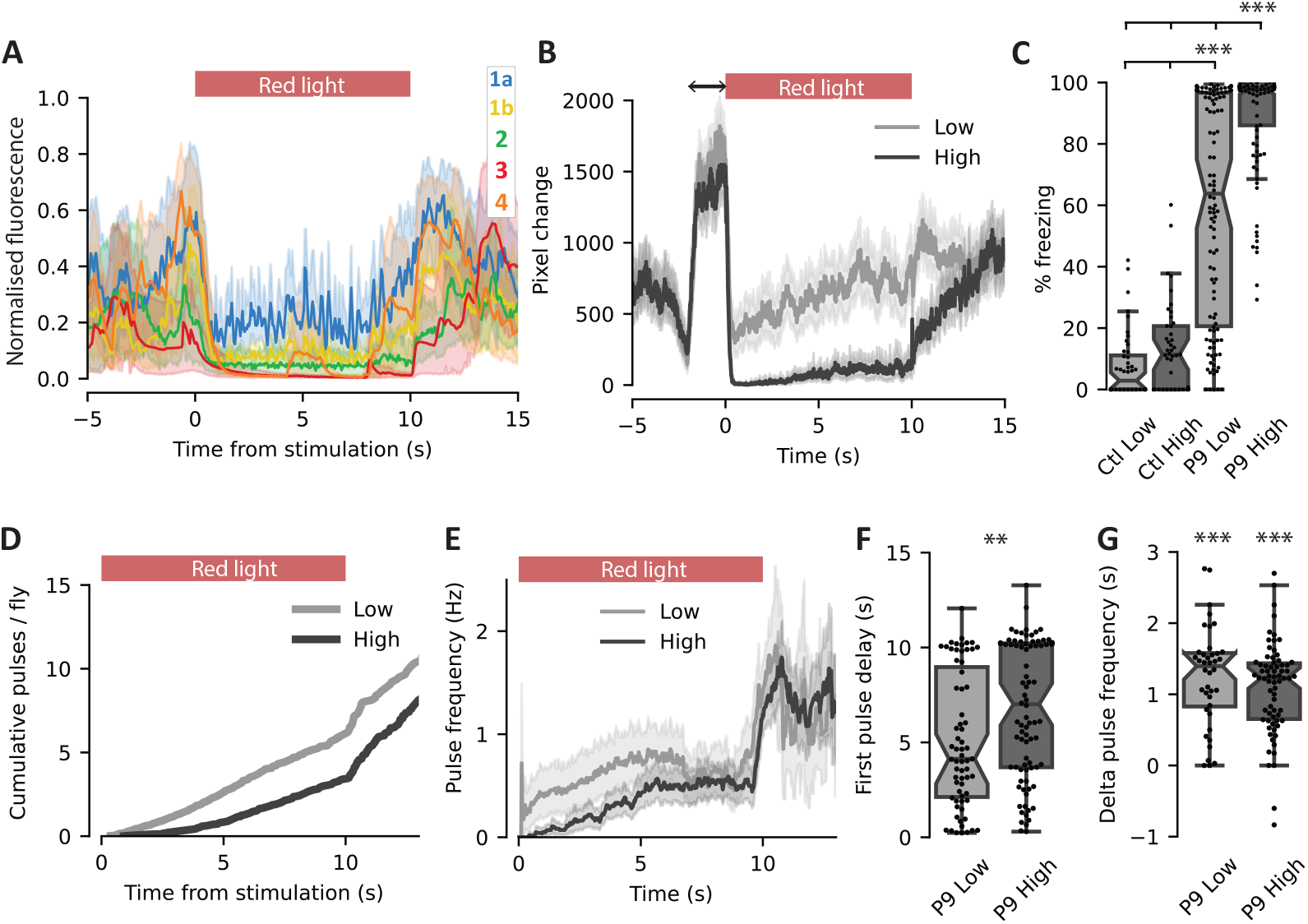
Freezing circuitry modulates pulse state. A) Average normalised muscle fluorescence for ROIs described in Figure 2, during optogenetic activation of DNp09 neurons in spinning disc microscope. Red bar indicates the time of stimulation. *N* = 9 flies. B) Average pixel change during DNp09 stimulation, for a ‘Low’ intensity stimulation (8.8 mW/cm^2^) and a ‘High’ intensity stimulation (21.5 mW/cm^2^). Black arrow indicates the 2 second window of required movement for the closed loop stimulation to be triggered. Red bar indicates stimulation time. *N* DNp09-Low = 114 stimulations (15 flies), *N* DNp09-High = 100 stimulations (13 flies). C) Percentage freezing during the 10 s stimulation. *N* Control-Low = 44 stimulations (6 flies), *N* Control-High = 43 stimulations (6 flies), *N* DNp09-Low = 114 stimulations (15 flies), *N* DNp09-High = 100 stimulations (13 flies). *P* DNp09 High < 0.001 compared to all other groups. PDNp09 Low < 0.001 compared to both control groups (Dunn’s test). D) Cumulative pulses per sample over the stimulation period for DNp09 Low and High flies. *N* DNp09-Low = 114 stimulations, *N* DNp09-High = 100 stimulations. E) Pulse frequency during and following the stimulation period, for DNp09 Low and High conditions. *N* DNp09-Low = 100 stimulations, *N* DNp09-High = 114 stimulations. F) Delay between freezing onset and first pulse. *N* DNp09-Low = 67 stimulations, *N* DNp09-High = 86 stimulations. *P* = 0.0015 (Mann Whitney U test). G) Pulse frequency difference between the latter half of the stimulation (5-10 s) and the 2 seconds after stimulation offset, for DNp09 Low and High conditions. *N* DNp09-Low = 40 stimulations, *N* DNp09-High = 70 stimulations. Statistics indicate the difference from 0. *P* < 0.001 for both conditions (Wilcoxon signed-rank test).

We next investigated the effect of DNp09 stimulation intensity on freezing level and pulsing state on our tethered set-up. We hypothesised that increased activation of DNp09 would lead to a deeper freezing state, as indicated by reduced pulsing. To control for any effect of light alone we used flies with the same genetic make-up but lacking the DNp09 enhancer (Empty-Gal4). We used two stimulation light strengths: 8.8 mW/cm^2^ (‘Low’) and 21.5 mW/cm^2^ (‘High’), presented as 10 seconds of constant light, delivered when the fly was engaged in walking behaviour (see Methods). As hypothesised, DNp09-activated flies froze upon light presentation and the level of freezing shown scaled with the light intensity, with the ‘Low’ flies showing lower levels of freezing (Fig. 5B and C, *P* < 0.001), reinforcing that levels of DNp09 activity are instrumental in determining the fly motor behaviour. Control flies showed negligible levels of freezing to the red light (Fig. 5C, Fig. S5B, *P* < 0.001). We next examined muscle pulsing during light-induced freezing. At both stimulation strengths, we observed both Continuous and Delayed phenotypes (Fig. S5C and D), and in both cases the pulse frequency ramped up throughout the freezing bout (Fig. S5E), phenocopying pulse dynamics during spontaneous or threat-induced immobility. During the stimulation, pulsing was decreased in the High intensity as compared to the Low (Fig. 5D and E), and the latency to initiate pulsing was significantly longer in this condition (Fig. 5F, *P* = 0.0015), suggesting that, in line with our prediction, stronger stimulation shifts the flies to a more delayed, deeper freezing state. Intriguingly, a striking increase in pulse frequency was observed upon red-light offset in both conditions (Fig. 5E and G, *P* < 0.001), suggesting that after DNp09 stimulation is released, the enduring freezing state shifts closer to movement resumption. These results indicate that DNp09 activity induces immobility during the freezing response by suppressing leg muscle contraction in a graded manner. At lower levels of activation it silences leg muscles but spares the accessory hearts’ pulsing, and when strongly active it also suppresses the hearts, potentially leading to a deeper state of freezing.

We also leveraged DNp09 activation to ensure that indeed an absence of calcium activity in muscles was equivalent to a non-contracted state. We expressed Chrimson in DNp09 neurons of flies expressing the tropomyosin-GFP fusion as described in Fig. 2. Sarcomeres, the contractile units of striated muscle, could be clearly visualised in the spinning disc microscope (Fig. S5F). Upon DNp09 stimulation, the distance between Z discs (lines delineating the end of each sarcomere unit) in the femur of the leg showed a significant increase as compared to timepoints without a stimulus occurring (Fig. S5G, *P* = 0.0104), indicating a relaxation of the muscles, and validating our previous hypothesis that the drop in calcium observed upon DNp09 stimulation was indeed representative of muscle relaxation.

### Induced accessory heart activation reduces freezing

Finally, we asked what controls the activity of the tibial accessory heart. We identified a line from the Janelia FlyLight collection, 92A06, whose anatomy revealed a single motor neuron per leg showing ramifications in the distal tibia (Fig. 6A), as well as sparse labelling in the brain and VNC (Fig. S6A). The soma and dendritic projections of the 92A06 motor neuron closely resembled those of the motor neuron described as targeting the tarsus flexor muscle in a recently published study mapping motor neuron projections in the Ventral Nerve Cord (VNC) to their muscular targets (37). A 3D reconstruction of the axonal projections of 92A06 showed neuro-muscular junctions (NMJs) targeting the 1a (pulsing) muscle fibres, as well as NMJs extending up the tarsus flexor muscle into the 1b fibres (Fig. 6B, Supp Video 8). This anatomy makes the 92A06 motor neuron an excellent candidate to carry the signal for the two types of activity in fibres 1a and 1b described in Fig. 2 – one which activates just the distal fibres in a pulsing manner (ROI 1a), and one which engages the pulsing muscle and the remaining fibres of the tarsus flexor muscle (ROI 1b) in a strong sustained contraction, which could be achieved by different synaptic strengths in the NMJs of the different muscle fibres.

**Fig. 6.**
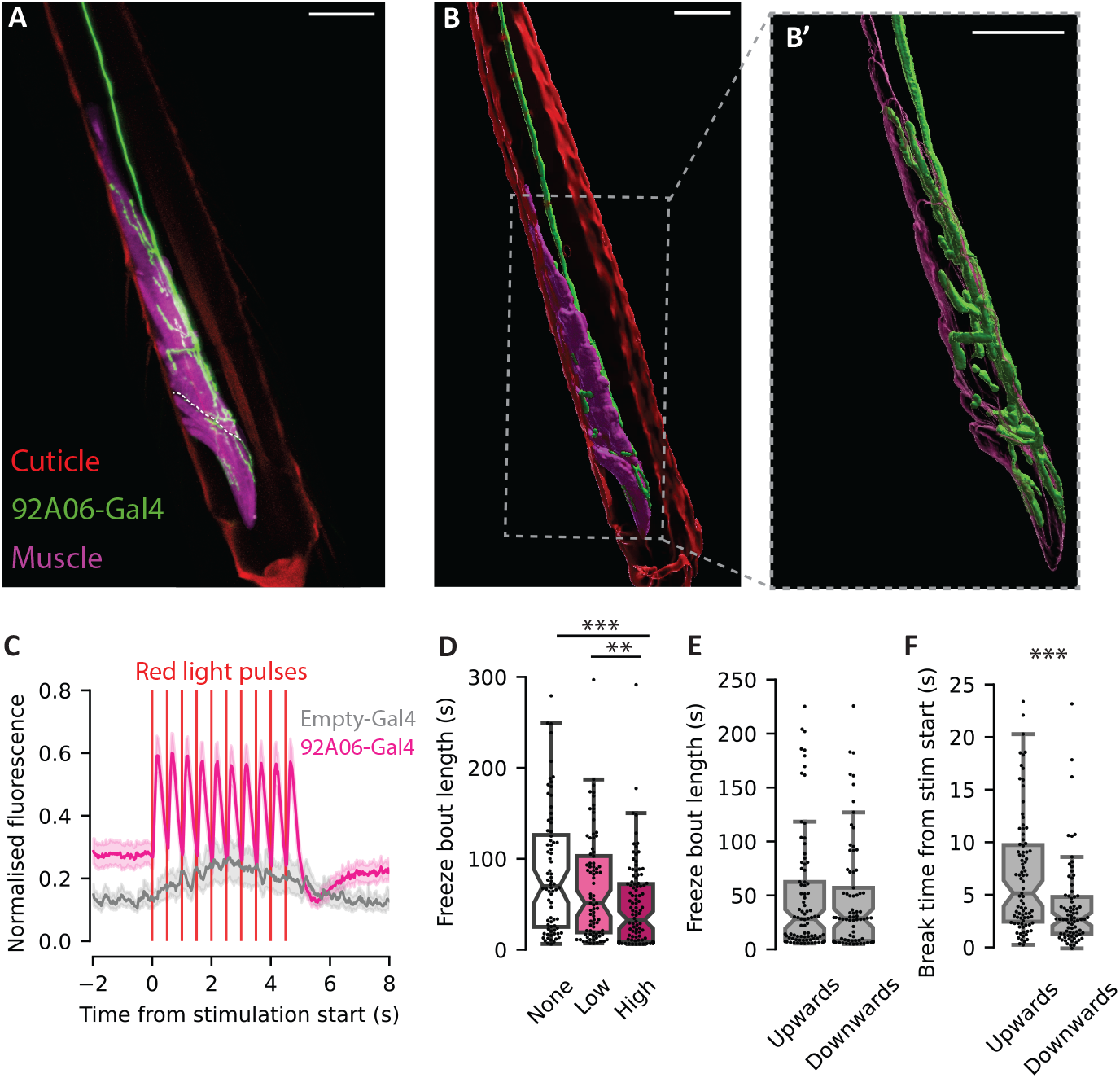
Induced pulsing causes freezing breaks. A) Confocal projection of the distal tibia. Red: cuticle autofluorescence. Green: Tibial ramifications of 92A06 line (labelled with GFP). Magenta: Tarsus flexor muscle (labelled with mhc-RFP, manually segmented). The white dotted line indicates the separation between the pulsing fibres (lower) and the rest of the tarsus flexor muscle (upper). Scale bar = 50 µm. B) 3D reconstruction of 92A06 motor neuron and tarsal flexor muscle. Colours as in (A). Inset: zoom-in on area indicated by grey box, showing the post-synaptic boutons of the 92A06 motor neuron entering the muscle. Scale bar = 50 µm. C) Fluorescence of 1a muscle fibres during activation of 92A06 motor neuron with Chrimson, and of Empty-Gal4 control. Red vertical lines show the times of 100ms pulses of red light. *N* Empty-Gal4 = 57 bouts (5 flies). *N* 92A06-Gal4 = 63 bouts (5 flies). D) Freeze bout lengths after loom presentation in 92A06-Gal4 flies, during the delivery of 3 different Chrimson stimulation frequencies: High (2.5 Hz), Low (0.5 Hz) and None (no light). *N* High = 108 bouts (30 flies). *N* Low = 81 bouts (26 flies). *N* None = 76 bouts (27 flies). *P* High-Low = 0.004, *P* High-None < 0.001, *P* Low-None = 0.52. (Dunn’s test). E) Freeze bout lengths after loom presentation, during delivery of either an upwards ramping or a downwards ramping stimulation. *N* Upwards = 87 bouts (23 flies). *N* Downwards = 77 bouts (22 flies). *P* = 0.66 (Mann-Whitney-U test). F) Moment when the fly broke from freezing during the final stimulus train of the freezing bout, for upwards and downwards ramping stimulations. *N* Upwards = 87 bouts (23 flies). *N* Downwards = 77 bouts (22 flies). *P* < 0.001 (Mann-Whitney-U test).

To confirm that the 92A06 motor neuron activated the pulsing muscle, we used optogenetic activation of the neuron with Chrimson. Expression of Chrimson in 92A06 neurons and GCaMP6f in muscles partially disrupted pulsing activity even when red light was not actively being shone on flies, whilst in control flies expressing the Empty-Gal4 promoter pulsing remained intact (Fig. S6B and C). Chrimson is known to have leaky activity (47) so it is possible that low uncoordinated activation impairs endogenous pulsing, suggesting that indeed these motor neurons are engaged in the pulsing activity of the tibial accessory heart. Light stimulation of tethered flies (10 pulses of 100 ms at 2 Hz) revealed a striking activation of the muscle fibres of the tarsus flexor muscle (Fig. 6C). To exclude the possibility that any of the off-target cells in the brain or VNC could be responsible for the muscle activation we observed upon optogenetic stimulation, we severed the leg from the body, thus removing all labelled cells except the motor neuron axon from the preparation, and imaged the leg on the spinning disc microscope while delivering the red-light stimulation. We observed clear muscle activation caused by the red light, confirming that activation of only the motor neuron axon was sufficient to cause activation of the tarsus flexor muscle (Fig. S6D).

We sought to understand how manipulating pulsing via changes to the activity of the 92A06 neurons could affect fly behaviour. We first attempted to eliminate pulsing; however, we were unable to achieve a full silencing, making an interpretation of the results difficult (Fig. S6E). Instead, we tested how artificial induction of pulsing (occurring in addition to the residual endogenous pulsing activity) changed freezing dynamics. Flies were tethered and exposed to looming stimuli, using the same experimental protocol as in Figures 1 and 2. When flies froze in response to a loom, we delivered red-light stimulus trains to cause muscle pulsing. We first tested whether inducing extra pulsing activity in the legs could cause early breaks from freezing. We delivered red light in 100 ms pulses at a constant frequency, at either 2.5 Hz or 0.5 Hz, as well as a no-light control. Empty-Gal4 control flies did not show any significant differences in freezing time to the different stimulation levels, showing that light delivery alone did not change freezing times (Fig. S6F). However, for flies expressing Chrimson under the 92A06 promoter, stimulation at a high frequency reduced freezing times compared to those receiving either low stimulation or no light (Fig. 6D), demonstrating that indeed the pulsing of the muscle can be causal for hastening a freezing break. to

We previously observed that the frequency of the endogenous pulsing ramped up to movement onset (Fig. 3B and C, Fig. S5F). We thus tested whether ramping stimuli, mimicking naturally occurring pulsing preceding movement onset, affected the time of breaking from freezing. We designed two stimulus protocols; an ‘upwards ramping’ stimulus resembling pulse dynamics during naturalistic freezing, and a ‘downwards ramping stimulus’ which consisted of the same inter-pulse intervals but reversed. If the flies remained freezing after the stimulus, it was re-delivered 10 seconds later. We observed that the muscle pulsing could accurately follow the changes in pulse frequency (Fig. S6G and H). We then analysed the freezing bout length for both conditions. We found that both types of ramping stimuli overall caused freezing breaks to the same degree (Fig. 6E). We next analysed whether the dynamics of the moment of breaking from freezing were the same between groups. Interestingly, we found that the flies did not break at the same moment for different stimuli. Flies given the ‘Downwards’ stimulation, which received high frequency stimulation at the beginning of the stimulus train, showed shorter latencies to break from freezing than flies receiving the ‘Upwards’ stimulation, which received the high frequencies later in the train (Fig. 6F). These data suggest that it is the high frequency pulsing that most strongly facilitates movement resumption, rather than the ramping pattern. Interestingly, in a small but notable percentage (20.6%) of the breaks for ‘Upwards’, the flies did not break during the stimulation itself but in the 10 seconds after the stimulation, suggesting that the stimulation may have induced a preparedness state which lead to subsequent movement. This was almost absent for the ‘Downwards’ case (3.8%) (Fig. S6I). The endogenous ramping of pulsing frequencies towards movement onset may reflect a progressive process of preparedness to move, suggesting it results from the integration of information over time, such as social cues discussed above. Together, these data show that physiological signals from the body, such as accessory heart pulses, are not simply reflections of an internal state but can act as feedback signals to the brain to promote an action.

## Discussion

In this work, we set out to interrogate the active and dynamic nature of freezing, a sustained defensive immobility state, through a combination of whole-body skeletal muscle imaging, anatomical reconstruction, and behavioural and neuronal manipulations. We describe a newly discovered rhythmic pulsatile muscle activity present in the distal tibia of *Drosophila*, which we reveal as a previously unidentified fly cardiac organ a leg accessory heart. The leg hearts’ activity was unique to immobility, both threat-induced freezing and spontaneous pausing, and was perfectly correlated between all the legs of the fly. Pulsing invariably started before the onset of whole-body movement, and pulse rate ramped up to movement onset. Higher threat levels led to less pulsing and a predominance of a Delayed phenotype over Continuous pulsing, while a virtual ‘safety’ cue caused an upregulation of pulse frequency and a higher probability of movement resumption. Finally, we identified a motor neuron that innervates the leg accessory heart and whose activation induced pulsing leading to premature breaks from freezing. In short, this study describes the anatomy and function of a hitherto unidentified accessory leg heart with pulsatile activity that is highly regulated, dynamically preparing flies for movement. Importantly, it reveals that threat-triggered freezing can be both a quiescent, likely more passive state, and an active preparatory state, that changes as a function of the external environment.

### Pulsing acts as an accessory leg heart

We describe a specialisation of the 3 distalmost fibres of the *Drosophila* tarsus flexor muscle, which pulse rhythmically during immobility, compressing a haemolymph-filled cavity at the end of the tibia. This compression could force the flow of haemolymph effectively through the leg, aiding circulation, thus performing the role of an accessory heart. Insect circulatory systems are open, in contrast to the closed systems of vertebrates, meaning the haemolymph flows in close contact with tissues without using discrete vessels. Low pressure in these systems presents a problem for adequate circulation, even with the help of a central dorsal heart. The situation is complicated further by the insect body organisation characterised by long thin limbs with high energy requirements, enclosed by a rigid exoskeleton which hinders the entry of haemolymph necessary to deliver nutrients to tissues. An evolutionary strategy used by insects to overcome this problem is a range of specialised pulsatile structures known as accessory hearts, which are found associated with a variety of appendages, strategically positioned to aid haemolymph supply, and act autonomously from the main dorsal heart (reviewed in (40)). The most widely studied of these is the wing heart, described in almost all insect orders, reflecting the critical role of flying for the success of most insects (reviewed in (48)). Antennal hearts are also extremely widespread and show great diversity amongst species. Other pulsatile organs appear to be less common and may reflect specialisations of anatomy or ecology in different species (49, 50).

Leg hearts were the first accessory pulsatile organ to be discovered, due to rhythmic pulsations of the tibiae observed in the aquatic *Hemiptera Notonecta glauca* (51), described in (39). Despite this extensive history, and the long length and narrow diameter of legs being impediments for haemolymph flow, until the present study there were reports of leg hearts in only the locust and in Hemiptera (39, 52). The structure and location of the heart in these two taxa is very different, with the locust displaying two specialised muscles in the trochanter and Hemiptera a muscle in the proximal tibia, suggesting an independent evolution. The muscle pulsing of the distal tarsus flexor fibres in *Drosophila melanogaster* we describe in this work, whilst differing in anatomical location to the previously described leg hearts, is highly congruent with them in terms of perceived function through pressure exerted on haemolymph in the adjacent enlarged sinus. The fibres retain their connection to the tarsus flexor tendon and are also active during overt leg movements. This would explain why pulsing was apparently limited to immobility, since during motion as this muscle contracted the fly would receive a boost to circulation ‘for free’, akin to the contribution of skeletal muscle activity to effective venous blood return in vertebrates or the pauses observed in the larval *Drosophila* heart during crawling (53). The co-option of a part of a canonical muscle for a new pulsing role is comparable to the leg heart in Hemiptera, where a subset of fibres of the pretarsus flexor muscle have been recruited to perform the accessory heart function (39).

The discovery of a new type of accessory heart in *Drosophila* shows the extraordinary variety of evolutionarily convergent solutions displayed in insects to the problem of appendage haemolymph flow. As well as enriching our knowledge of the basic physiology and anatomy of *Drosophila*, our study sheds light on the central control and coordination of the peripheral hearts. Whilst it has been known since the first description of leg hearts that their control and pumping frequency is independent of the dorsal heart (51), and that leg hearts could have bouts of pumping and quiescence (39), there were no descriptions of pumping patterns or their relations to the behaviour of the animal. Through extensive behavioural studies we show for the first time the modulation of accessory heart pulsatile activity in accordance with the state of freezing and the readiness to move. The remarkable synchrony of the tibial pulsing muscles between all leg suggests a centrally controlled neuronal circuit independent of the locomotor circuits that control leg joints, which are coupled in an out-of-phase fashion. Additional circuit analysis experiments, including the connectome definition of the upstream motor neurons, will shed light on to the underlying circuits that control leg pulsing activity and their modulatory features.

### Pulsing reveals changing states during freezing

Immobility is a characteristic response to a threat observed across the animal kingdom. It has been a subject of research for centuries and has been described by many different names throughout time; ‘freezing’, ‘tonic immobility’, ‘thanatosis’, ‘playing dead’, ‘animal hypnosis’, ‘death feigning’, ‘playing possum’, ‘catalepsy’ and more (21). This multiplicity hints at the diversity of states which could be concealed underneath the static exterior. At the two extremes, a ‘freezing’ state is usually considered a highly alert, reactive state seen at intermediate threat levels where the animal is assessing the situation and is poised for action, while ‘playing dead’ or ‘tonic immobility’ implies animal is unalert and practically unresponsive to external cues, for minutes to hours after the initial threatening or harmful stimulus (10). These states are very unlikely to be merely binary, but instead may encompass a spectrum of immobile behaviours between the two. The internal state, and the implications for the animal’s physiology and future behaviour, will vary widely across this spectrum.

To understand the behavioural control of threat-induced immobility more fully, we utilised our discovery that the *Drosophila* leg heart was highly modulated by the immobility state. The pulsing rhythm and frequency of the muscle was variable during freezing, even within the same fly (Fig. 3A, Supp Fig. 4C), revealing that the outwardly identical static exterior concealed a variety of internal and neural states. More intriguing still was our discovery that the pulsing frequency ramped up to the moment of movement onset, and that a voluntary, controlled freezing break was never observed without at least one preceding pulse. This led us to hypothesise that pulsing represented a ‘preparedness-to-move’ state, with at least one pulse being necessary before movement resumed. In line with this hypothesis, freezing bouts with extended delays before pulses started (‘Delayed’) were on average longer than those which had more steady frequencies (‘Continuous’). Pulsing alone was not sufficient to break freezing (some freezing bouts showed continuous pulsing for more than a minute), indicating there is an additional layer of control over the choice to break or not from freezing – however inducing extra pulses increased the likelihood of movement onset (Fig. 6). It is possible that pulsing reflects a holistic internal state progression with multiple layers of control leading to the resumption of movement.

These results strongly indicate that the immobility seen during freezing can encompass a multiplicity of states. A continuously pulsing fly may be eminently poised to exit freezing during the whole bout but will stay in place until a ‘go’ signal (likely from a higher brain centre) is received, whereas others enter a deeper state of quiescence and may progress to the pulsing state and then to the freezing break almost simultaneously.

### Pulsing as a ‘preparedness-to-move’ state

Empirical reports of extended muscle engagement during immobility to physiologically prepare for movement are scarce. Some insects such as bees have been shown to use skeletal muscles as thermoregulators to increase the thoracic temperature in preparation for flight (54). However, the present study is the first that we are aware of in insects to demonstrate causal relationship between the activity of a cardiac muscle during immobility and the preparation to resume movement. Movement is very costly due to the high energy requirements of muscle contraction; thus, preparatory physiological changes to avoid shortages of nutrients or oxygen needed for aerobic respiration upon the onset of movement make conceptual sense. (Note that while the insect haemolymph does not transport oxygen directly, internal fluid pressure changes can play a respiratory role by augmenting gas flow through the tracheal system (29).) However, experimental reports of such mechanisms are sparse. There is evidence from vertebrates that respiratory upregulation can occur immediately prior to commencement of movement (55–57). Gravel et al. (2007) showed that in lampreys, spontaneous swimming activity was sometimes preceded by a long burst of respiratory brain activity occurring up to 8 seconds before movement started. In humans, mental simulation of exercise without actual movement causes a modest increase in both heart rate and ventilation rate (55, 58). Although direct evidence of similar mechanisms in invertebrates is lacking, it has been shown that during a prolonged tonic immobility state the Colorado potato beetle displays regular abdominal pumping movements, which have a demonstrated respiratory function in various insect species (59), during which the metabolic rate doubles, suggesting that these movements could represent an analogous ‘movement preparation’ state (60). It is interesting to consider that the nervous system and the physiological output of the accessory heart appear to have a reciprocal interaction, whereby nervous activation can alter pumping characteristics (Fig. 5), but muscle activation can also lead to a behavioural change such as breaking from freezing (Fig. 6).

It would be interesting to see whether the pulsing state and preparation for movement signatures that we observe in freezing hold true for other immobile states such as sleep. Recently, proboscis pumping to aid in waste clearance by upregulating the flow of haemolymph was described during fly sleep (61). If the goal of this deep sleep state is to increase haemolymph flow, it would make sense that other pumping mechanisms such as accessory hearts were activated at the same time, and indeed there is a report of correlated tarsal twitches during sleep, resembling the twitches we report in Figure 2 caused by contraction of the pulsing muscle (44).

Our work opens new avenues for the study of brain-body interactions during defensive behaviour, which have recently started to yield fascinating results on the interconnectivity of body and affect (62), and of how these correspond to adaptive internal states. More broadly, it contributes to our understanding of how immobility is achieved, maintained and finally broken, deepening our understanding of the multi-stages process involved in the preparation for movement, with wide implications for the mechanisms of defensive freezing and action planning in general.

## Methods

### Reagents, fly stocks and softwares

See Key Resources table in Supplementary Material for full details.

### Fly lines and husbandry

All flies used in this study were non-virgin females *Drosophila melanogaster* between 3 and 7 days post-eclosion. Flies were housed in a 25° incubator with 50-70% humidity and a 12h:12h light:dark cycle, and reared on standard food, except for flies used for optogenetics. These flies were fed food supplemented with all-trans-retinal (0.4mM, Sigma-Aldrich), starting at least 2 days before experiments started, and kept in the dark until being used for experiments. The 92A06-Gal4 line was generated *de novo* by plasmid injection (a gift from Todd Laverty) using standard procedures in the attP2 landing site.

### Experimental procedures

#### Tethering procedure

Flies were briefly cold anaesthetised. A small blob of Norland 81 optical adhesive on a custom-made wire holder was lowered onto the fly’s thorax and cured with 365 nm UV lamp (UVP) for 30s on each side of the fly. Flies were given 10 minutes to recover inside the behavioural apparatus before the experiment started. To glue a single leg (experiments in Fig. 1F-M), after the gluing of the thorax, Norland 81 optical adhesive was applied to the vertical section of the wire, and the left hind leg was gently stretched upwards into the glue. The UV lamp was used for 30s to cure the hind leg glue. Given that manipulation of a single leg risks causing damage to its fragile structure during process of gluing, we took as a ground truth our result that activity should be correlated between the glued leg and the free legs during moments of immobility (Fig. 1D and E) and excluded flies which did not meet this criterion, i.e. showing an average correlation of below 0.4 (threshold based on previous analysis), as having likely suffered lesions to leg muscles and neurons. For analyses of glued leg fluorescence shown in Figure 1E-M, only bouts of movement or stillness of more than 2 seconds were considered. Peak widths and peak ITIs are plotted excluding data outside the 99th centile.

#### Tethered Behavioural apparatus

The tethering wire was positioned over a small porexpam ball (4.5 mm in diameter), held in a custom-made 3D printed holder. To reduce reflection, the ball was coloured black at least a day before experiments using a layer of paint (Posca, uni) and ink (Sharpie). Flies could freely pick up the ball, move it and release it. To excite GCaMP6f, flies were illuminated with a blue LED (470 nm, Thorlabs) with a blue filter (488 nm, Thorlabs). To image movement, flies were illuminated with an infra-red LED (850 nm, Thorlabs). A near-IR hot mirror (ThorLabs) was placed laterally to the fly to separate the light paths from the two different wavelengths of light. Green fluorescence was captured using a Hamamatsu C11440 camera (Orca-flash4.0LT) with Zeiss 50mm Makro-Planar lens and a green filter (510 nm, Thorlabs) at 50 fps using HC Image Live software. Infra-red light was captured using a FLIR Flea3 camera with a Computar macro-zoom lens and a long-pass filter (800mn) at a frame rate of 120 fps using custom written Bonsai workflows (which also controlled any optogenetic stimulations). To synchronise the two imaging paths, a counter signal was delivered to the imaging capture softwares every 5s in the form of either a TTL pulse or a photodiode signal. The rig was placed inside an opaque black box.

#### Visual stimuli

Visual stimuli were presented to the fly using an LG 24” monitor with a refresh rate of 144 Hz, positioned 22.5 cm from the fly and tilted at an angle of 30º. The monitor background was purple to remove luminance change artifacts in the fluorescence channel during looming stimuli. As described in the text, a vertical black stripe, known as a buridan stripe (31, 63) with a width of 23º on the fly’s visual field was presented on the monitor throughout the entirety of the experiments to increase walking behaviour, in all tethered fly results except those in Figure 1F-M and Figure 4C-J. Buridan stripe opacity could be either 100% (resulting in a 0% contrast difference with the loom) or 80% (20% contrast difference with the loom), as specified in the text. Looming stimuli were generated using custom-written scripts in PsychoPy, based on formulae from (64). Looming stimuli consisted of a black circle expanding on the background. The visual angle of the expanding circle was determined by: *q*(*t*) = 2*tan*^*−*1^(*l/vt*), where l is half of the length of the object and v the speed of the object toward the fly. Virtual object length was 1 cm and speed was 25 cm/s (l / v 40 ms). Each looming presentation lasted for 500 ms. The time of loom onset was signalled to both imaging streams using a photodiode triggered by the appearance of a small black square in the corner of the monitor. The first loom was presented after a baseline period of 5 minutes. For all experiments except those in Figure 4KM, subsequent looms were manually triggered once the fly had broken from freezing. If the fly did not freeze to a loom, there was a minimum inter-loom interval of 10 seconds before the next loom was presented. For experiments in Figure 4K-M, 20 looms were presented on a stereotyped schedule with an average inter-loom interval of 15s. The social movement stimulus in Figure 4C-J was generated by playing a video of 4 flies walking in a rectangular area on the stimulation monitor. Flies had an average length of 6.9 cm on the monitor (excluding the wings), giving a final visual angle of 17.2º to the focal fly. The video had a purple background as for other visual stimuli. The video was played throughout the 5-minute baseline of the experiment. At every loom presentation, the video initially paused during the 0.5s taken for the loom to expand and fill the screen. When the loom disc disappeared, the video reset to a still image of the initial frame (ensuring that the cue was the same for every trial), to give the impression of the virtual flies freezing, for a period of 10.7s. This timepoint was chosen empirically as sufficient to allow an even split between flies which showed no pulsing and flies which showed pulsing before the cue onset. If the focal fly was still freezing at this point, the ‘Cue’ was triggered. For the flies in the ‘Motion’ group, the video started playing to deliver the social motion cue to the focal fly. In the ‘No Motion’ group the video remained paused throughout the whole freezing bout of the focal fly and was manually triggered to restart when the focal fly broke from freezing. In both groups, when the focal fly broke from freezing there was a 15s inter-trial interval during which the video was playing before the next loom was delivered.

#### Optogenetic stimuli (in tethered set-up)

For optogenetic activation experiments in Figures 5 and 6, a deep red LED (wavelength 627 nm) positioned 3 cm away from the fly was used to activate CsChrimson. For experiments in Figure 5, light power at the fly was 8.8 mW/cm^2^ for ‘Low’ and 21.5 mW/cm^2^ for ‘High’ powers. Mounting and recovery were as described above. Flies were imaged for a 2 minute baseline, after which stimulation was triggered when the fly was walking, using a closed loop movement detection system in Bonsai (65) which required the fly to be walking for at least 80% of a 2 second window. Each stimulation consisted of constant red light for 10 seconds, followed by a minimum inter-stimulus interval of 40 s before light could be re-triggered by the movement of the fly. Each fly received multiple presentations (between 5 and 8) at the same stimulus strength. For all experiments in Figure 6, light power was 21.5 mW/cm^2^ at the fly, and stimuli had a duration of 100 ms. For experiments in Figure 6D, flies were imaged for a 5-minute baseline, after which a loom was delivered to flies to induce freezing. If the fly froze, a train of LED activation was delivered after 5 s. The train consisted of either a ‘High’ frequency stimulus (red light pulses of 100 ms delivered at 2.5 Hz); a ‘Low’ frequency stimulus (same as high but with a frequency of 0.5 Hz); or ‘None’ (same as high but with LED power set to 0). All stimuli were delivered in an unbroken train until the fly broke from freezing. After a freezing break, there was an inter-trial-interval of around 35 s before a new loom was delivered to the fly. For experiments in Figure 6E-F, baseline and loom delivery was as above. Light trains were delivered after 5 s, which could either be an ‘Upwards ramping’ stimulus, consisting of 10 red-light pulses delivered at 1.5 Hz, followed by 6 pulses at 2 Hz and 4 pulses at 2.5 Hz, or a ‘Downwards ramping’ stimulus of 4 pulses at 2.5 Hz, 6 pulses at 2 Hz and 10 pulses at 1.5 Hz. If the fly remained freezing for 10 seconds after the pulse train, another pulse train was delivered. If the fly broke from freezing, there was a minimum interval of 35 s before another loom was manually triggered. If the fly did not freeze at all upon the loom, a subsequent loom would be presented after 35 s.

#### Leg, VNC and brain dissection and mounting

Leg dissection was performed as described in (66). Briefly, legs were dissected from the body in PBS. The legs were then fixed in 4% PFA overnight at 4°C. After fixation legs were washed 3x in 0.3% Triton-X in PBS for 20 minutes at R.T. and mounted in Vectashield (VectorLaboratories). Brains and VNCs were fixed in 4% PFA for 20 minutes and dissected in PBS. Tissues were labelled with primary antibodies raised against brp (mouse nc82, 1:10 DSBH) and GFP (rabbit anti-GFP, 1:1000 ThermoFisherScientific), followed by secondaries Alexa Fluor 594 anti-mouse (1:500 ThermoFisherScientific) and Alexa Fluor 288 anti-rabbit (1:500 ThermoFisherScientific) respectively. Tissues were mounted in in Vectashield (VectorLaboratories).

#### Spinning disc and confocal imaging

Spinning disc experiments were performed as described in (67). Briefly, flies were cold anaesthetised for 2-3 minutes. Left legs were glued to a glass coverslip using UV curable glue. The thorax and abdomen of the fly were immobilised using beeswax. Imaging was performed on a Nikon/Andor Revolution XD spinning-disk confocal microscope with an EMCCD camera (iXon 897) using iQ 3.5 software and either a 10x dry objective or a 40x oil objective. Images were taken at a scan speed of 17-20 fps. For optogenetic stimulation experiments, a set of 6 deep red LEDs was paced around the fly to provide red light stimulation. Stimulation protocols and a 5 s TTL timing pulse were controlled by an Arduino. For severed leg experiments in Figure S6B, intact flies were glued as above. Flies were imaged, and 10 stimulus pulses delivered (100 ms, 2 Hz) to ensure muscles were visible. Without moving the leg embedded in the glue, the body and the leg were separated using forceps. The coverslip was returned to the microscope and 2-4 stimulus trains (10 pulses, 100 ms, 2 Hz, minimum inter-train-interval 20 s) were delivered starting when the muscle was in a low-contraction state. Fluorescence in the severed leg was normalised to maximum and minimum values from the intact fly video. Confocal imaging was performed on a Zeiss (Oberkochen, Germany) LSM980 confocal microscope with a 10x dry or 40x glycerol objective. Cuticle and tendons were captured without labelling due to strong autofluorescence in red and far-red channels (36).

#### Erioglaucine injections

Erioglaucine (Sigma-Aldrich) was diluted in PBS to a final concentration of 5 mg/ml. The solution was filtered to remove particles. Flies were cold anaethetised and 100 nl of solution was injected into the thorax, using pulled glass pipettes connected to a Nanoject 3 (Drummond). Flies were left to recover for 10-15 minutes. Flies were then glued to a coverslip using UV curable glue (Norland 81) and imaged in a Zeiss Axioimager M2 microscope.

### Data processing and analysis

#### Leg muscle nomenclature

Previous descriptions of *Drosophila* leg muscles have differed in the names used to describe antagonistic muscles which control leg segments (36, 37) (see (37) supplementary information for a discussion of the differences), with a combination of the terms ‘flexor/extensor’ and ‘levator/depressor’ in usage for different muscle pairings. For the ease of comprehension to a general audience, we have opted to use only the ‘flexor/extensor’ nomenclature.

#### Freezing identification

For tethered fly experiments, the infra-red video was used to identify moments of freezing defined as bouts of more than 100 ms where pixel change, as detected by a custom written Bonsai script, fell below a threshold of 90 pixels per frame (determined by manual annotation), and which could only be broken by movement bouts of more than 25 ms. The equivalent frames were extracted from the fluorescence videos for fluorescence processing. Immobility bouts where the fly dropped the ball were discarded. There were 2 bouts (out of 113) where the freezing classifier was broken without any preceding pulses – upon examination the fly was found to do a small movement (in one case with a single leg, in one case a body postural adjustment) and then resume freezing.

#### Fluorescence processing

*Tethered flies:* A custom-written ImageJ macro was used for extraction of pulsing muscle fluorescence during freezing bouts. ROIs were manually drawn around pulsing muscles for the duration of the freezing bout. For all experiments except those specified below, the single clearest pulsing muscle of the left legs (ipsilateral to the camera) was selected, i.e. one that did not show large changes in position and was not obscured by the body or wings. For experiments in Figure 1D and E, freezing segments where 4-6 of the pulsing leg muscles (both ipsilateral and contralateral to the camera) were clearly visible were selected. ROIs, consisting of the pulsing muscles plus 3 other regions for comparison, were manually drawn. For experiments in Figure S6B, C and E, 3 ROIs were drawn around the pulsing muscle of the three left legs of the flies. Mean fluorescence traces of the ROI were extracted for each freezing bout. Peak detection was performed in python, manually adjusting the detection parameters for every bout. For quantification of average pulse frequency, pulse moments were smoothed with a rolling filter of 51 frames.

*Immobilised flies:* Spinning disc fluorescence levels were extracted in ImageJ using manually drawn ROIs in 5 different areas of muscle. All fluorescence levels were normalised to the maximum and minimum values for individual muscle, for each sample.

#### Quantification and statistical analysis

All data analysis and statistical tests were performed using custom-written scripts for Python v3.10.9. Since the majority of data sets failed normality tests, we used non-parametric statistics throughout the study. Statistical tests are specified in figure legends. All tests are two-tailed. Multiple comparisons were adjusted using a Bonferroni correction. Violin plots features represent: centre line, median; upper and lower lines, 75th and 25th quartiles. Box plot features represent: centre notch, median; box limits, 75th and 25th quartiles; whiskers, 1.5x interquartile range. Line plots with variance represent (unless otherwise specified in the legend): line, mean; variance, 95% confidence interval. Statistics representation on figures: * signifies ≥0.05 *P*>0.01. ** signifies ≥0.01 *P*>0.001. *** signifies 0.001 ≥*P*.

#### Muscle activity state classifier (in Fig. 2D)

All muscle fluorescence values were normalised to fall between 0 and 1 as described above. To classify muscle activity states, we used a Butterworth high-pass filter with a cut-off of 2 Hz to identify ‘Pulsing’ states. For non-pulsing states, we smoothed the fluorescence signal using a rolling average with a 5-frame window. We defined ‘High’ states as those when the smoothed signal was above 0.2, and ‘Low’ states as those below 0.2. ‘Transition’ states were defined as states when the muscle was both pulsing and the average fluorescence was above 0.4 (these numbers were chosen using visual inspection of the traces).

#### Continuous/Delayed pulsing classifier (in Fig. 3)

To separate freezing bouts into Continuous and Delayed based on pulsing activity, we plotted the distribution of the percentage of all pulses found in the first 50% of the bout, for both the 0% contrast and 20% contrast data (from Figures 1 and 4). This gave us a bimodal distribution (Fig. S3A). We made a cut-off in the trough of the distribution, with bouts below 15% classified as Delayed and those above as Continuous. There were a few bouts which fell above this threshold but had extremely low pulse rates, which did not fit the conceptual framework of a ‘Continuous’ bout. We thus added a second criterion whereby continuous bouts needed to have a total average pulse rate of > 0.2 Hz.

#### Confocal image analysis

For leg images, ImageJ plug-ins for channel subtraction (to remove cuticle autofluorescence) and stitching of tiles were used (68, 69). 3D reconstructions (Fig. 2E, Supp Video 5 and Fig. 6B, Supp Video 8) were made using Imaris (Oxford Instruments). To distinguish and isolate individual muscle fibres, tendons and neuron, image stacks were manually segmented using ImageJ, before being processed in Imaris.

#### DeepLabCut tracking

DeepLabCut (43) was used to track 18 body points: 3 body points (head, thorax and tail) and 5 points per leg (Body-Coxa; Coxa-Femur; Femur-Tibia; Tibia-Tarsus; and Tarsus-end). For the group data only fore- and hind-leg tracking were used for the final analysis, since middle legs were frequently positioned perpendicular to the camera so joint angles could not be accurately calculated. Limb angles were calculated using python. Angles were normalised to always represent deflections in the downwards direction.

## ACKNOWLEDGEMENTS

We thank all members of the Moita lab, the Mendes lab and Rita Teodoro for their comments and insight throughout this project. We thank Natalia Barrios especially for proposing the idea of an accessory heart. We thank Rui Gonçalves, CONGENTO (Consortium for Genetically Traceable Organisms), the Champalimaud Fly Platform and the NMS Fly Platform for fly husbandry, Liliana Costa for dissections, Catarina Craveiro and Champalimaud MTTP platform for transgenic line generation, the Champalimaud Hardware platform, Artur Silva, Pedro Garcia and João Frazão for hardware assistance, and the Champalimaud ABBE platform, Anna Pezzarossa, Pedro Campinho and Telmo Pereira for imaging assistance. We thank Lalanti Venkatasubramanian, John Tuthill and Camile Guillermin for fly line suggestions, Todd Laverty at Janelia Research Campus for the 92A06 plasmid, and Bloomington Stock centre for fly lines. We thank Luisa Vasconcelos, Clara Ferreira, Natalia Barrios, Alisson Gontijo and Alfonso Renart for comments on the manuscript, and Gil Costa for illustrations. This work was supported by (ERCCoG819630-A-Fro) and (UIDB/04443/2020) to MAM, (PTDC/BIA-COM/0151/2020) and (UIDB/04462/2020, UIDP/04462/2020) to CSM, the research infrastructure CONGENTO (LISBOA-01-0145-FEDER-022170), and ABBE imaging (LISBOA-01-0145-FEDER-022122). AMM and CAR were supported by FCT doctoral fellowships, respectively (PD/BD/128445/2017) and (2023.03812.BD).

## Author contributions

AFH, CSM and MAM conceived and designed the study. AFH performed experiments and analysed data. MCF, AMM and CAR performed experiments and data processing for the following figures: MCF (Fig. 4 and 6), AMM (Fig. 2 and 5), CAR (Fig. 4 and 5). AFH, CSM and MAM wrote the manuscript. All authors contributed to discussions about the work and manuscript revision, and read and approved the submitted version. Correspondence and requests for materials should be addressed to Dr César Mendes or Dr Marta Moita

## Notes

### Competing Interest Statement

The authors have declared no competing interest.

