## Supplementary Info and Figures for "Covert muscle activity reveals dynamic freezing states and prepares the animal for action"

### Key Resources Table

| Reagent or resource | Source | Identifier |
|---|---|---|
| **Antibodies** | | |
| Mouse anti-nc82 | DSHB | RRID: AB_2314866 |
| Rabbit anti-GFP | ThermoFisherScientific | A-11122 |
| Alexa Fluor 488 anti-rabbit | ThermoFisherScientific | A-11034 |
| Alexa Fluor 594 anti-mouse | ThermoFisherScientific | A-11032 |
| **Chemicals** | | |
| Norland 81 optical adhesive | Norland Products | https://norlandprod.com/ |
| Erioglaucine | Sigma-Aldrich | 861146 |
| All-trans-Retinal | Sigma-Aldrich | R2500 |
| **Experimental models: *D. melanogaster*** | | |
| Canton-S | | BDSC_64349 |
| mhc-LexA,LexAOp-GCaMP6f | (1) | |
| CLIP190-Gal4 | (2) | BDSC_65638 |
| tropomyosin[cc00578] GFP | | BDSC_51537 |
| Tin-CGal4 | (3) | |
| UAS-CsChrimson | (4) | BDSC_55139 |
| 92A06-Gal4 | This paper | |
| DNp09-Gal4 | (5) | BDSC_75903 |
| mhc-RFP | | BDSC_38464 |
| 10XUAS-IVS-eGFPKir2.1 | (6) | |
| UAS-CD8::GFP | | BDSC_32185 |
| **Software and algorithms** | | |
| Python v3.10.9 | Anaconda | https://www.anaconda.com/ |
| ImageJ (Fiji) | (7) | https://fiji.sc/ |
| Bonsai v2.6.3 | (8) | https://open-ephys.org/bonsai |
| HCImageLive | Hamamatsu | https://hcimage.com/hcimage-overview/hcimage-live/ |
| Imaris 10.2.0 | Oxford Instruments | https://imaris.oxinst.com/ |
| PsychoPy v2020.2.5 | (9) | https://www.psychopy.org/ |
| DeepLabCut | (10) | https://www.mackenziemathislab.org/deeplabcut |

### Supplementary Video legends

Supplementary Video 1

***Drosophila melanogaster* show coordinated leg pulsing of a distal tibia muscle during freezing**

First video: fly shows Continuous pulsing (see Figure 3D). Second video: fly shows Delayed pulsing (Figure 3E).

Supplementary Video 2

**Freely moving flies show leg pulsing during freezing and grooming**

Freely moving flies expressing GCamP6f in muscles were filmed in a small behavioural area. Flies showed both freezing to a loom (first video, loom not shown) and grooming behaviour (second video). Correlated activity of distal tibial muscles could be observed.

Supplementary Video 3

**Muscle activity in an immobilised leg**

Spinning disc confocal microscopy time-lapse of the immobilised leg of a fly expressing GCaMP6f in the muscles. Muscle labels refer to those described in Figure 2a. The outline of the tibia and femur is indicated by the grey dashed line. 20 fps.

Supplementary Video 4

**Muscle 1a and Muscle 1b show a semi-dissociation of activity**

Animated graph showing the relationship between fluorescence in Muscle 1a and Muscle 1b over time. Fluorescence for both muscles was normalised to extend between 0 and 1.

Supplementary Video 5

**3D reconstruction of *Drosophila melanogaster* distal tibia**

3D reconstruction of the distal tibia, showing the locations of the cuticle (red), tendons (white) and muscles (green, magenta, blue and cyan). The fibres of the pulsing muscle are distinguished from the rest of muscle 1a by their colours.

Supplementary Video 6

**Shape changes of Muscle 1a as it contracts**

Time-lapse imaging of labelled muscles (genotype: tropomyosin GFP fusion) during contraction. The fibres of muscle 1a were manually annotated with a red line.

Supplementary Video 7

**Muscle 1a activity and tarsus twitching in unbraced legs**

Videos show simultaneous recording from the infrared signal and fluorescence signal, in flies expressing GCaMP6f in the muscles. Video 1: Fly with all legs off the ball. Video 2: Fly with only one leg, the medial leg ipsilateral to the camera, off the ball.

Supplementary Video 8

**3D reconstruction of 92A06 motor neuron, and Muscle fibres 1a and 1b.**

3D reconstruction of the distal tibia, showing the locations of the cuticle (red), tarsal flexor muscle (magenta) and 92A06 motor neuron (green).

### Supplementary Figures

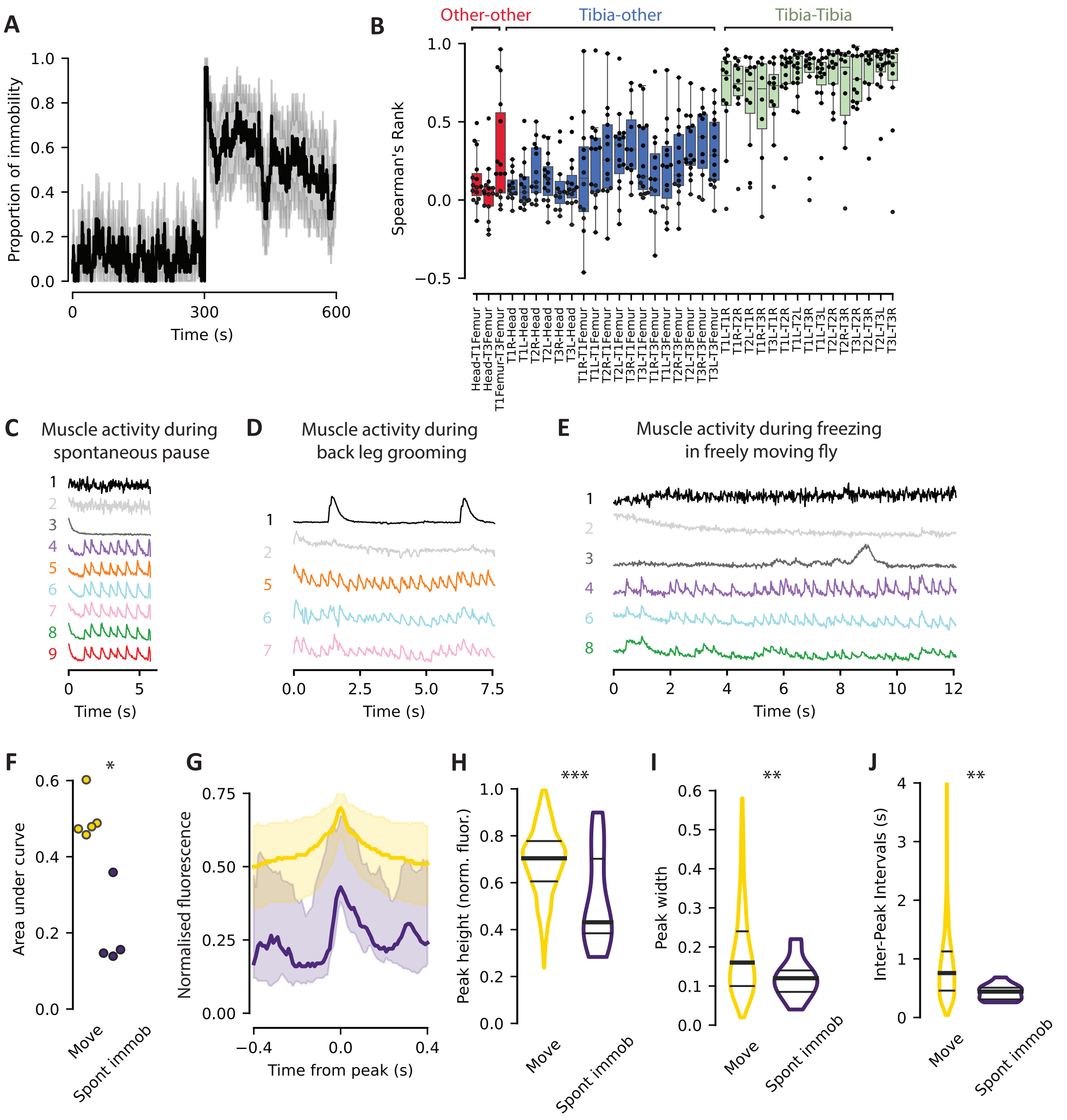

**Fig. Supp 1.** A) Overall proportion of freezing throughout the experiment. $N$ = 26 flies. B) Spearman's correlation values for 36 pairwise comparisons between the 9 ROI regions in Figure 1C. $N$ = 15 bouts (11 flies). C) Individual example of fluorescence in the ROIs during a spontaneous pause during the baseline period. ROI colours are consistent with those in Figure 1C. D) Individual example of pulsing during back leg grooming in 5 visible and stable ROIs. ROI colours are consistent with those in Figure 1C. E) Individual example of pulsing during freezing in a freely moving fly, in 6 visible ROIs. ROI colours are consistent with those in Figure 1C. Baseline fluctuations seen in ROI 8 are due to abdomen muscles captured in the ROI. F) Integration of area under fluorescence intensity curve per second, for movement vs freezing bouts during the stimulation period. $N$ = 5 flies. $P$ = 0.016, Mann Whitney U test. G) Average peak shape (median and interquartile range) for movement vs spontaneous immobility during baseline period. $N$ Movement = 1498 peaks, $N$ Spontaneous immobility = 22 peaks (5 flies). H) Normalised height of fluorescence peaks during baseline period. $N$ Movement = 1500 peaks, $N$ Spontaneous immobility = 23 peaks (5 flies). $P < 0.001$, Mann Whitney U test. I) Width of fluorescence peaks during baseline period. $N$ Movement = 1500 peaks, $N$ Spontaneous immobility = 23 peaks (5 flies). $P$ = 0.008, Mann Whitney U test. J) Inter-peak intervals during baseline period. $N$ Movement = 1479 IPIs, $N$ Spontaneous immobility = 12 IPIs (5 flies), $P$ = 0.008, Mann Whitney U test.

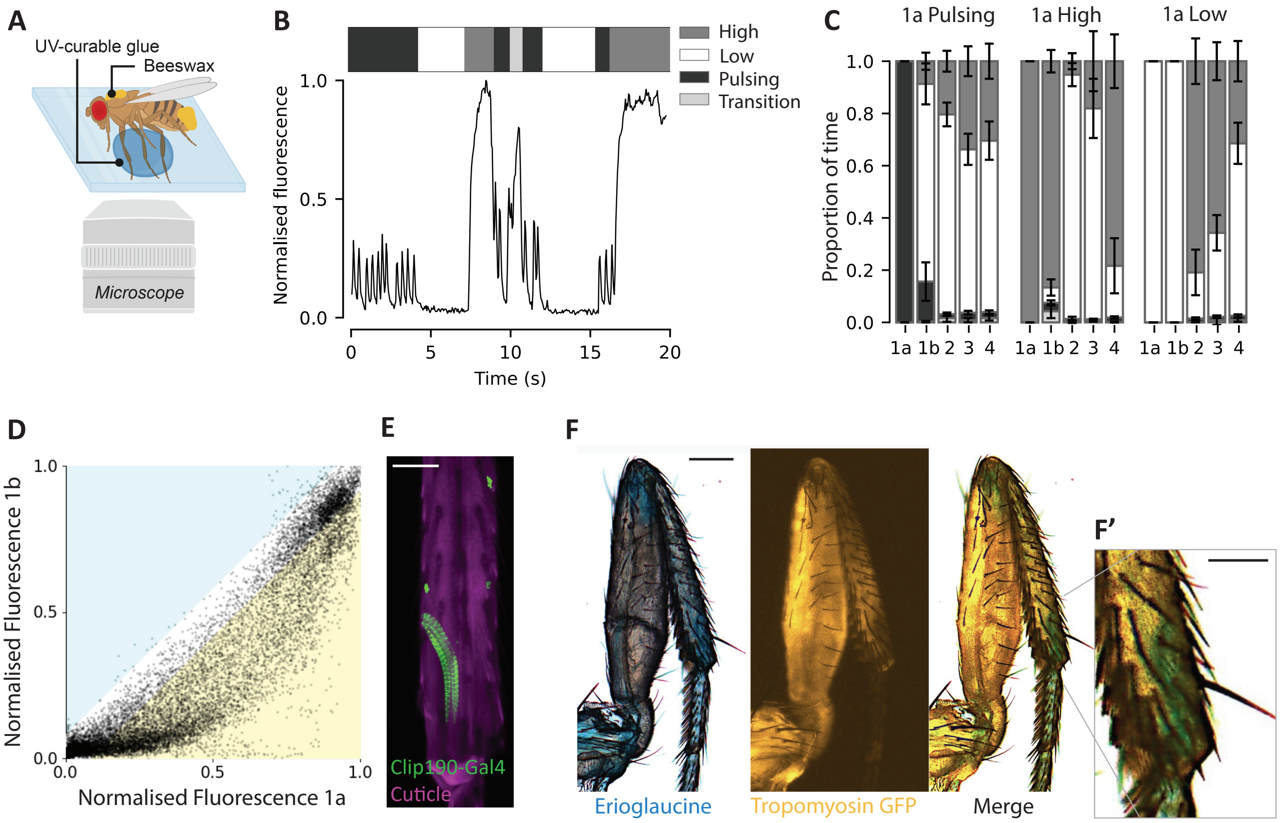

**Fig. Supp 2.** A) Schematic of spinning-disc confocal set-up. B) Example of muscle activity classified into 4 activity states (High, Low, Transition and Pulsing). The normalised fluorescence trace is shown below, with the classified states above. C) Proportion of time spent in the classified states, for all the time muscle 1a was in a Pulsing state (left); in a High sustained state (middle); or a Low state (right). $N$ = 9 flies. Error bars = SEM. D) Moment-to-moment correlation of normalised fluorescence for muscles 1a and 1b. $N$ = 9 flies. E) Confocal image of the Clip-180-Gal4 line expressing GFP. Scale bar = 40 µm. F) Distribution of haemolymph stained with erioglaucine, and muscles (tropomyosin-GFP). Scale bar = 100 µm. Inset: zoom-in of the indicated section of the distal tibia. Scale bar = 50 µm.

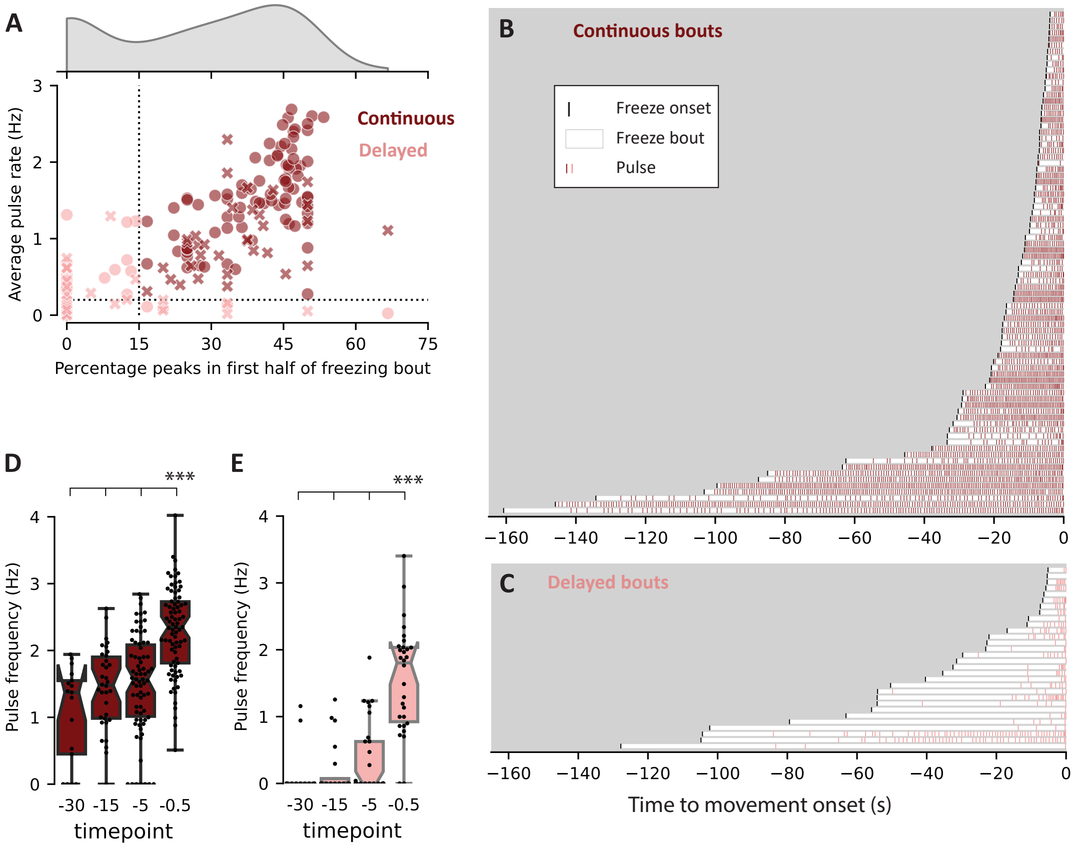

**Fig. Supp 3.** A) Cut-off boundaries for Continuous and Delayed bouts (See Methods for full description and reasoning). Dots: data from experiments with Low Saliency loom (Figs 1, 3, 4A and 4B). Crosses: data from experiments High Saliency loom (Fig. 4A and 4B). $N$ = 140 bouts, 45 flies. B) Continuous freezing bouts longer than 4 seconds, showing moments of muscle pulsing. Bouts are aligned to movement onset and ordered by length of freezing bout. $N$ = 81 bouts. C) As in (B) for Delayed bouts. $N$ = 30 bouts. D) Indicated timepoints of pulse frequencies for Continuous bouts. $N$ = 17 (-30s), 34 (-15s), 74 (-5s), 81 (-0.5s), $P$ = 0.001 (Nemenyi-Friedman test). E) As in (D) for Delayed bouts. $N$ = 16 (-30s), 20 (-15s), 30 (-5s), 30 (-0.5s), $P$ = 0.001 (Nemenyi-Friedman test).

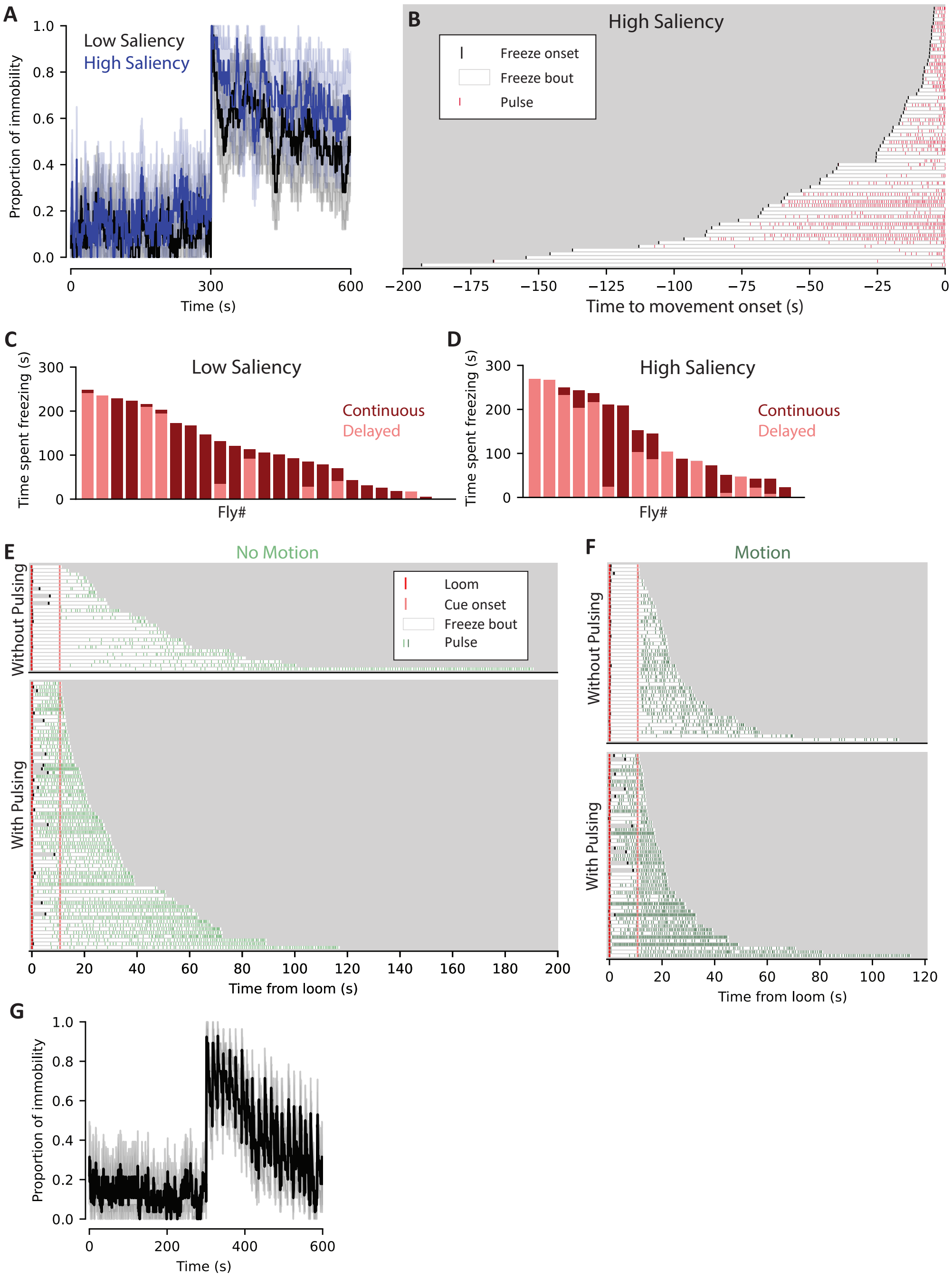

**Fig. Supp 4.** A) Overall proportion of immobility throughout the experiment for Low Saliency (same data as in Figure 1 and 3) and High Saliency. *N* Low Saliency = 25 flies, *N* High Saliency = 20 flies. B) Freezing bouts longer than 4 seconds, showing moments of muscle pulsing, for High Saliency looms. Bouts are aligned to movement onset and ordered by length of freezing bout. *N* = 71 bouts. C) Total amount of time spent freezing in either Continuous or Delayed mode, for Low Saliency looms. *N* = 25 flies. D) Same as in C, but for High Saliency looms. *N* = 20 flies. E) Freezing bouts longer than 4 seconds, showing moments of muscle pulsing, for flies which receiving the No Motion cue. Data is divided into 'Without Pulsing' (no pulses before cue onset) and 'With Pulsing' (at least one pulse before cue onset). Bouts are aligned to the loom presentation and ordered by time of movement onset. *N* Without pulsing = 28 bouts, *N* With pulsing = 65 bouts (33 flies). F) As in E but for flies receiving the Motion cue. *N* Without pulsing = 45 bouts, *N* With pulsing = 53 bouts (35 flies). G) Overall proportion of immobility throughout the Auto-loom experiment (cropped at 600s), *N* = 28 flies.

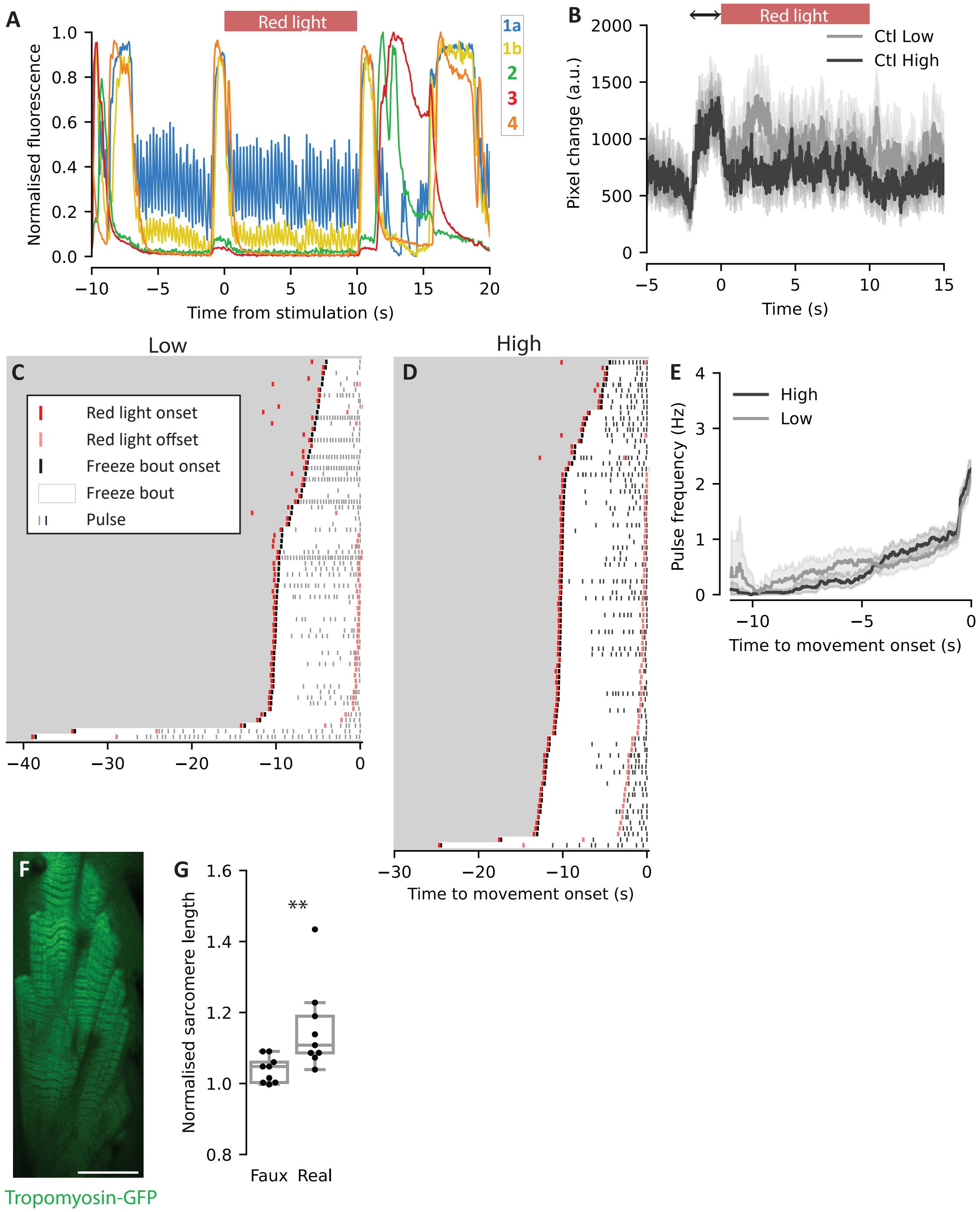

**Fig. Supp 5.** A) Individual example of immobilised leg muscles during optogenetic stimulation of DNp09 neurons in the spinning disc set-up. ROIs are as described in Figure 2. B) Average pixel change during Empty-Gal4 stimulation, for a 'Low' intensity stimulation (8.8 mW/cm$^2$) and a 'High' intensity stimulation (21.5 mW/cm$^2$). Black arrow indicates the 2s window of required movement for the closed loop stimulation to be triggered. Red bar indicates stimulation time. *N* Control-Low = 44 stimulations (6 flies). *N* Control-High = 43 stimulations (6 flies). C) Freezing bouts following DNp09 stimulation for 'Low' intensity stimulation, aligned to the moment of movement onset and ordered by length of freezing bout. *N* = 68 bouts. D) As in C but for 'High' intensity stimulation. *N* = 87 bouts. E). Pulse frequency during freezing bouts leading up to movement onset. *N* Low = 68 bouts (15 flies), *N* High = 87 (13 flies). F) Spinning-disc confocal image of femur flexor muscle expressing Tropomyosin-GFP, oriented with Proximal at the top and Distal at the bottom. Scale bar = 30 µm. G) Length of sarcomeres following a sham stimulation or a real stimulation of p9 neurons. *N* = 9 stimulations (5 flies). *P* = 0.01 (Mann Whitney U test).

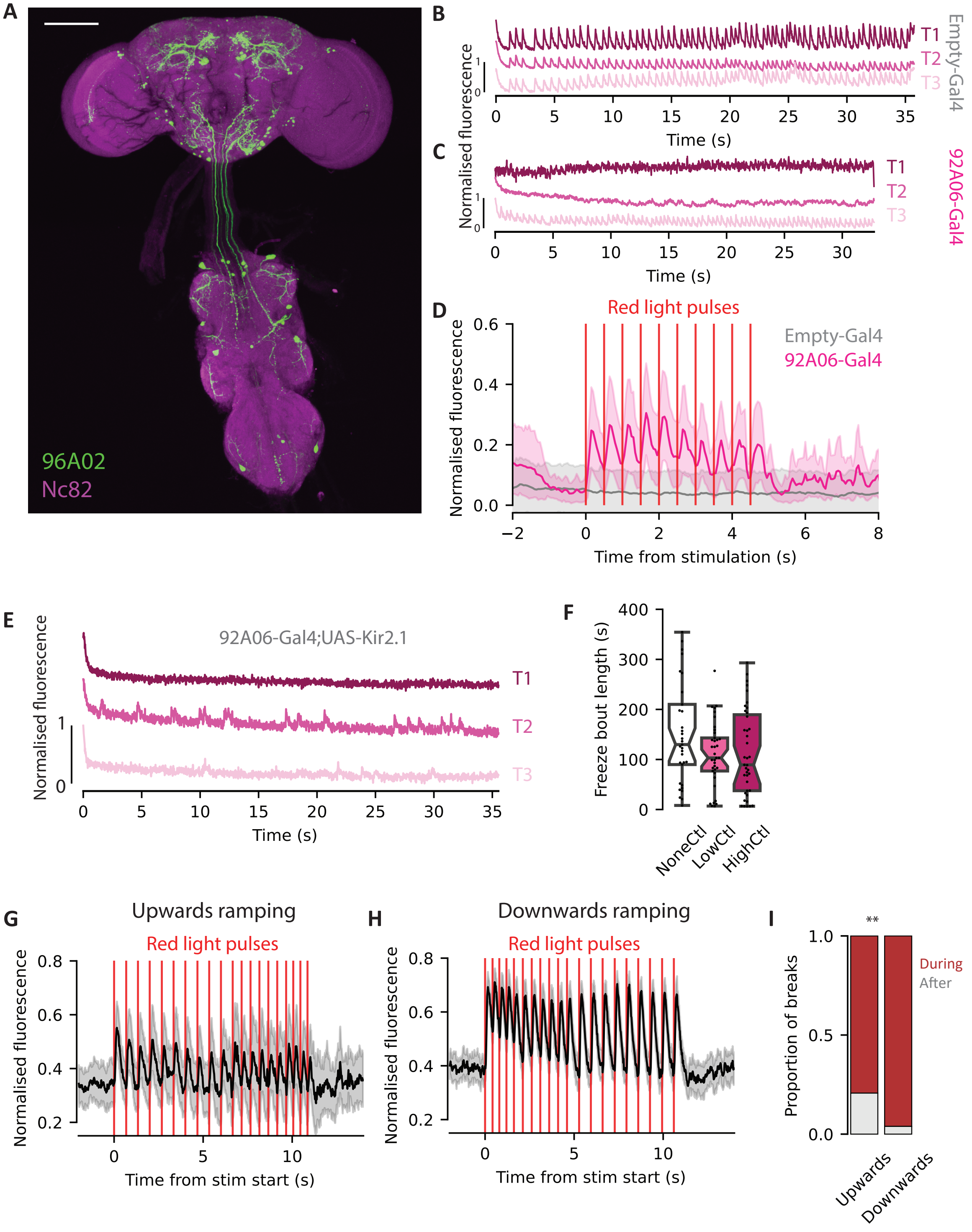

**Fig. Supp 6.** A) Brain and VNC projections for 92A06 line. Scale bar = 100 µm. B) Example fluorescence changes in pulsing muscle fibres (1a) in 3 legs during freezing in a control fly (Empty Gal4). C) As in (B) for 92A06-Gal4. D) Fluorescence changes in pulsing fibres (1a) during stimulation of the 92A06 line, in a leg severed from the body. *N* Empty-Gal4 = 9 stimulations (3 flies). *N* 92A09-Gal4 = 14 stimulations (6 flies). E) Example fluorescence changes in pulsing fibres during freezing in a fly expressing hyperpolarising channel Kir2.1, driven by 92A06-Gal4. F) Freeze bout lengths after loom presentation in Empty-Gal4 flies, during the delivery of 3 different Chrimson stimulation frequencies: High (2.5 Hz), Low (0.5 Hz) and None (no light). *N* High = 41 bouts (16 flies). *N* Low = 37 bouts (16 flies). *N* None = 33 bouts (16 flies). *P* High-Low = 1, *P* High-None = 0.27, *P* Low-None = 0.38. (Dunn's test). G) Fluorescence traces in pulsing muscle fibres during an upwards ramping stimulation. H) As in G for a Downwards ramping stimulation. I) Proportion of freezing breaks that happen during or after the stimulation train, for the upwards and downwards ramping cases. *N* Upwards = 87 stimulations (23 flies). *N* Downwards = 77 stimulations (22 flies). *P* = 0.003 (Chi-squared test).
